# Stereotyped goal-directed manifold dynamics in the insular cortex

**DOI:** 10.1101/2023.11.13.566809

**Authors:** Itay Talpir, Yoav Livneh

## Abstract

The insular cortex is involved in diverse processes including bodily homeostasis, emotions, and cognition. Yet we lack a comprehensive understanding of how it processes information at the level of neuronal populations. We leveraged recent advances in unsupervised machine learning to study insular cortex population activity patterns (i.e., neuronal manifold) in mice performing goal-directed behaviors. We find that the insular cortex activity manifold is remarkably consistent across different animals and under different motivational states. Activity dynamics within the neuronal manifold are highly stereotyped during rewarded trials, enabling robust prediction of single-trial outcomes across different mice, and across various natural and artificial motivational states. Comparing goal-directed behavior with self-paced free consumption, we find that the stereotyped activity patterns reflect task-dependent goal-directed reward anticipation, and not licking, taste, or positive valence. These findings reveal a core computation in insular cortex that could explain its involvement in pathologies involving aberrant motivations.

## Introduction

The insular cortex (InsCtx) is involved in a variety of processes, ranging from basic functions including bodily homeostasis, emotions, motivation, decision making, social interactions, and sensory processing, to maladaptive conditions including drug addiction, chronic pain, anxiety, obesity, and eating disorders^1–8^. Yet we lack a comprehensive mechanistic understanding of the computations in InsCtx that underlie its contribution to so many diverse processes.

A central path to understanding the computations within and across brain regions lies in measuring and deciphering neural activity of large populations of neurons in different behavioral contexts^9–11^. Indeed, there has been a recent explosion of artificial intelligence and machine learning approaches to decipher neural activity at the level of large neuronal populations^12–14^. These methods involve exploration of the multi-dimensional structure of activity patterns, also known as the activity manifold. By doing so, these approaches have begun to provide insights into the core computations of neural circuits, such as motor cortex control of limb movement, thalamic representation of head direction, and entorhinal cortex representation of spatial position^15–19^. Most previous animal model studies of InsCtx activity in behaving animals have used various recording and analysis techniques, including fiber photometry^20–23^, single neuron electrophysiology^24–31^, and ensemble analyses of electrophysiological recordings from relatively small populations (∼5-10 neurons)^32–37^. While these studies have substantially advanced our understanding of InsCtx, they have not fully leveraged these recent artificial intelligence and machine learning approaches. Therefore, there is potentially much to be gained from comprehensive unbiased investigations of activity patterns of large InsCtx populations, using these recent developments in neuroscience analytical tools.

Experimental investigation of the population activity manifold can be used to test predictions from theoretical studies making explicit assumptions regarding the precise computations in the network (e.g., aforementioned studies of head direction and spatial position^15–17,38–41)^. Alternatively, unbiased analyses of the manifold structure using, e.g., unsupervised machine learning, can aim to infer computations from the revealed manifold structure and dynamics within. Our study here falls within the latter category.

Previous work has suggested that InsCtx activity on short-timescales (∼seconds) represents multi-modal gustatory and interoceptive sensory stimuli^26,42–46^, as well as multi-modal salient external cues^24,25,31,36,44,47–50^. Further work has suggested that such activity may also convey emotional, motivational and valence information^20–22,51–53^. Longer timescale changes in InsCtx activity (minutes to hours) have been suggested to represent slow changes in the physiological state of the body^22,47,54–57^, as well as specific positive and negative emotional states^20–22,51^. However, seeing as InsCtx integrates many different inputs and is involved in many diverse processes, understanding its function at the neuronal population level requires comprehensive analyses of the repertoire of its activity patterns (i.e., the neuronal manifold) and their dynamics. Doing so would help answer fundamental questions that remain unanswered. For example, how constrained are InsCtx population activity patterns by intrinsic factors vs. extrinsic factors, such as the current behavioral context? How are different external and internal behavioral variables represented concomitantly in InsCtx population activity patterns? Are there core computations that InsCtx performs across different contexts? If so, will they have distinct signatures in population activity space? Here, we leverage recent advances in unsupervised machine learning and topological data analysis to perform an unbiased investigation of InsCtx population activity during goal-directed behaviors.

## Results

### Insular cortex population activity manifold structure is stereotyped across different mice and motivational states

To investigate how neuronal activity patterns are structured in InsCtx, we analyzed two-photon calcium imaging data from previously published and unpublished datasets from layer 2/3 of mid-InsCtx (**Table S1**). We started with data from thirsty (water-restricted) mice performing an operant go/no-go visual discrimination task in which licking following three initially arbitrary visual cues (drifting gratings) leads to rewarding (water), aversive (1M NaCl), or neutral outcomes^55^ (**Figure 1A-B**). We previously showed that most InsCtx neurons respond to aspects of this behavioral task with either increase or decrease in activity (e.g., visual cues, licking, reward consumption)^55^. Therefore, to detect any underlying manifold structure within the observed activity patterns, we compared the temporal similarities across all simultaneously recorded neurons. Importantly, points in the manifold represent population activity patterns at specific times (**Figure 1C-D**).

**Figure 1–.**
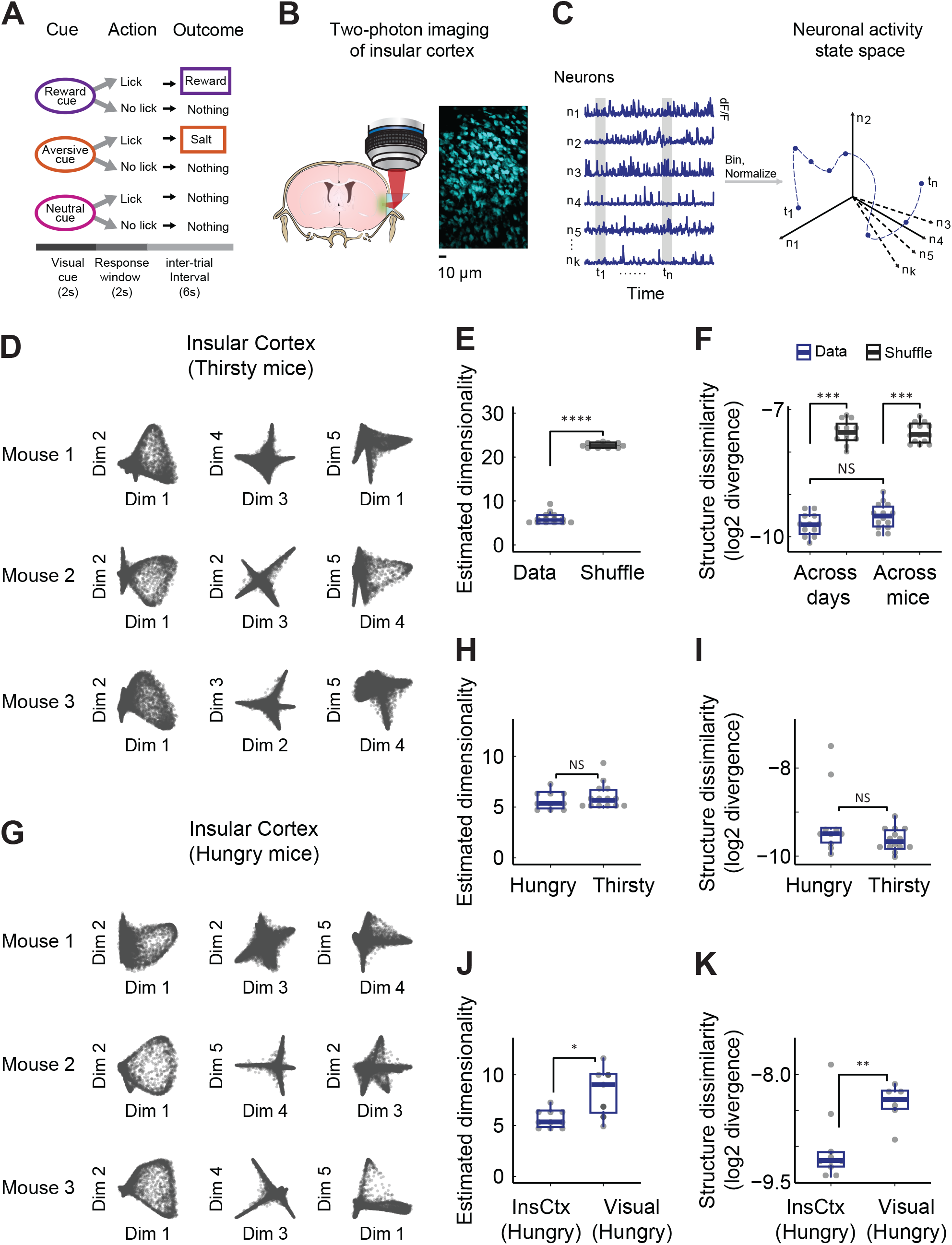
InsCtx activity manifold structure is stereotyped across different mice and motivational states. **A.** Schematic of the operant visual discrimination task. **B.** Two-photon calcium imaging of InsCtx through a microprism: schematic coronal brain section illustrating the approach, and example field-of-view. **C.** Illustration of the approach to study InsCtx population activity. **D.** Three representative planes of dimensionality-reduced neuronal activity over time, from three different thirsty mice. Each point represents the population activity pattern in a 0.5 sec time bin. Note the high qualitative similarity between different mice. **E.** Estimation of intrinsic dimensionality for each individual dataset vs. shuffled data. The estimated dimensionality was significantly lower than chance (P<7*10^−5^, one-tailed Wilcoxon signed-rank test). **F.** Quantification of topological dissimilarity (*β*0 features) between all datasets (see also **Figure S2**). Notably, there was a significant difference when comparing either across days or across mice to chance levels (P<0.0003, one-tailed Wilcoxon signed-rank test), while comparisons of the topological dissimilarity across days and across mice yielded no significant differences (P>0.1, two-tailed Wilcoxon rank sum test). **G.** Same as in ‘**D**’, for hungry mice. Different mice than in ‘**D**’. **H.** Similar estimated intrinsic dimensionality across hungry and thirsty mice (P>0.4, two-tailed Wilcoxon rank-sum test). **I.** Similar activity manifold shape (topological dissimilarity, *β*0 features) across hungry and thirsty mice (P>0.25, two-tailed Wilcoxon rank-sum test). **J.** Comparison of estimated intrinsic dimensionality in InsCtx and cortical visual areas (POR – dark shading, V1 – light shading) in hungry mice performing the same behavioral task. *P<0.015, one-tailed Wilcoxon rank-sum test. **K.** Comparison of activity manifold shape (topological dissimilarity, *β*0 features) between InsCtx and visual areas in hungry mice performing the same behavioral task. **P<0.0035, one-tailed Wilcoxon rank-sum test.

Previous studies have shown activity manifolds to be low dimensional during simple behavioral tasks, i.e., they can be described by relatively few variables as compared to the number of recorded neurons^10,13,15,16,41,58–60^. However, because InsCtx integrates multimodal information from somatosensory, auditory, visceral, gustatory, and other limbic regions^61,62^, we wondered whether InsCtx activity would be similarly low dimensional. Importantly, in all our analyses we analyzed each dataset in its entirety, including all time-point from all trials, starting from a thirsty (or hungry) state, and ending in a quenched (or sated) state. We estimated the dimensionality of InsCtx population data^58,63^ (**Figure S1A-B**), and found that across all datasets activity manifolds consistently had 6±1 dimensions (mean ± SD). This was independent of the specific parameters used to assess dimensionality and of the number of recorded neurons (**Figure 1E, S1C-E**). This consistently low intrinsic dimensionality, together with the diverse multi-modal inputs InsCtx receives, suggests substantial convergence and integration of information in InsCtx during goal-directed behavior.

Building on this finding, we used non-linear dimensionality reduction to unveil the low dimensional manifold structure of InsCtx activity^12,13,15–17,41,60,63^. Specifically, we used Laplacian Eigenmaps (LEM), due to its capability to effectively examine the local geometry of high-dimensional data^41,64,65^. This provided a finer-grain approximation, particularly beneficial when studying dynamics. Using LEM, we reduced dimensionality to the mean estimated intrinsic dimensionality across all datasets (6 dimensions). This uncovered a remarkably consistent activity manifold structure across datasets and different mice (**Figure 1D, S1F**). To quantify structure similarity, we used Topological Data Analysis, an emerging approach for quantitatively describing complex data structures^12,15,66–68^. We measured structure similarity by assessing the difference (i.e., divergence) between the distributions of topological features across datasets (see procedure illustration in **Figure S2A**). This revealed that the low-dimensional manifold structure remained consistent across different days and mice. Structural similarity was significantly lower for shuffled datasets (**Figure 1F**). We further validated these findings using pairwise permutation tests^69^ (**Figure S2B-C**).

We next assessed whether InsCtx dimensionality and manifold structure are conserved across different motivations. We compared data from thirsty (water-restricted) mice working for water rewards and hungry (food-restricted) mice working for liquid food rewards within the same behavioral task structure^50^ (**Figure 1G**). Activity manifold dimensionality in hungry and thirsty mice were quite similar (5.7±1 vs. 6±1; mean ± SD; **Figure 1H**). Moreover, activity manifold structure in hungry and thirsty mice were similar (**Figure 1I**).

We wondered whether the consistent dimensionality and topological similarity we observed in InsCtx across similar behaviors during different motivations merely reflect behavioral constraints on brain-wide activity^70–72^. We compared the InsCtx activity manifold with the activity manifold of visual cortical areas of mice performing the same behavioral task during the same hunger motivation (primary visual cortex and postrhinal cortex^73^). We found that the dimensionality and structure of the activity manifold in visual areas were both significantly higher and less consistent than those in InsCtx (**Figure 1J-K, S3**), and this did not depend on the number of sampled neurons (see Methods). We further confirmed this by examining the distribution of P-values from pairwise comparisons of manifold structure similarity between datasets within each group (**Figure S2D**). Notably, dimensionality and structure (topology) can be independent (e.g., two-dimensional datasets could exhibit either ring or sheet structures). As such, the fact that we observed different structures (topology) between InsCtx and visual areas, cannot be explained by the higher variability in the estimated dimensionality of the visual areas datasets.

Collectively, our findings show that the activity manifold in InsCtx (but not visual areas) is low-dimensional and structurally similar across mice engaged in similar goal-directed behaviors. The consistent InsCtx manifold structure across animals may indicate common encoded information. Moreover, these characteristics likely do not result from brain-wide constraints on behavior, and could thus be more specific to a subset of brain regions, including InsCtx

### Insular cortex activity dynamics within the manifold are stereotyped across different mice

Based on previous work^22,24,50,52,55^, we expected InsCx activity patterns on both short and long time-scales to correlate with external and internal behavioral variables (e.g., rewarding/aversive outcomes, satiety, arousal, etc.). We color-coded the manifold based on different experimental parameters (**Figure 2A**, left and middle; **S4A**) and found that some time-points associated with a given variable aggregated spatially within the activity manifold, while others were more uniformly distributed. We developed an automated method to cluster time-points within the manifold (termed here as “clustered structure”). This method clustered time-points that diverged from the center of the manifold (‘manifold clusters’), which was not assigned to any cluster (‘central point cloud’; **Figure 2A**, right; **S4B-C**). Notably, we consistently found 7 manifold clusters in 46/47 of analyzed datasets. Note that different cluster indices (represented by different colors; **Figure 2A**) were randomly assigned during clustering, and therefore do not necessarily map onto the same behavioral variables across datasets (see more below).

**Figure 2–.**
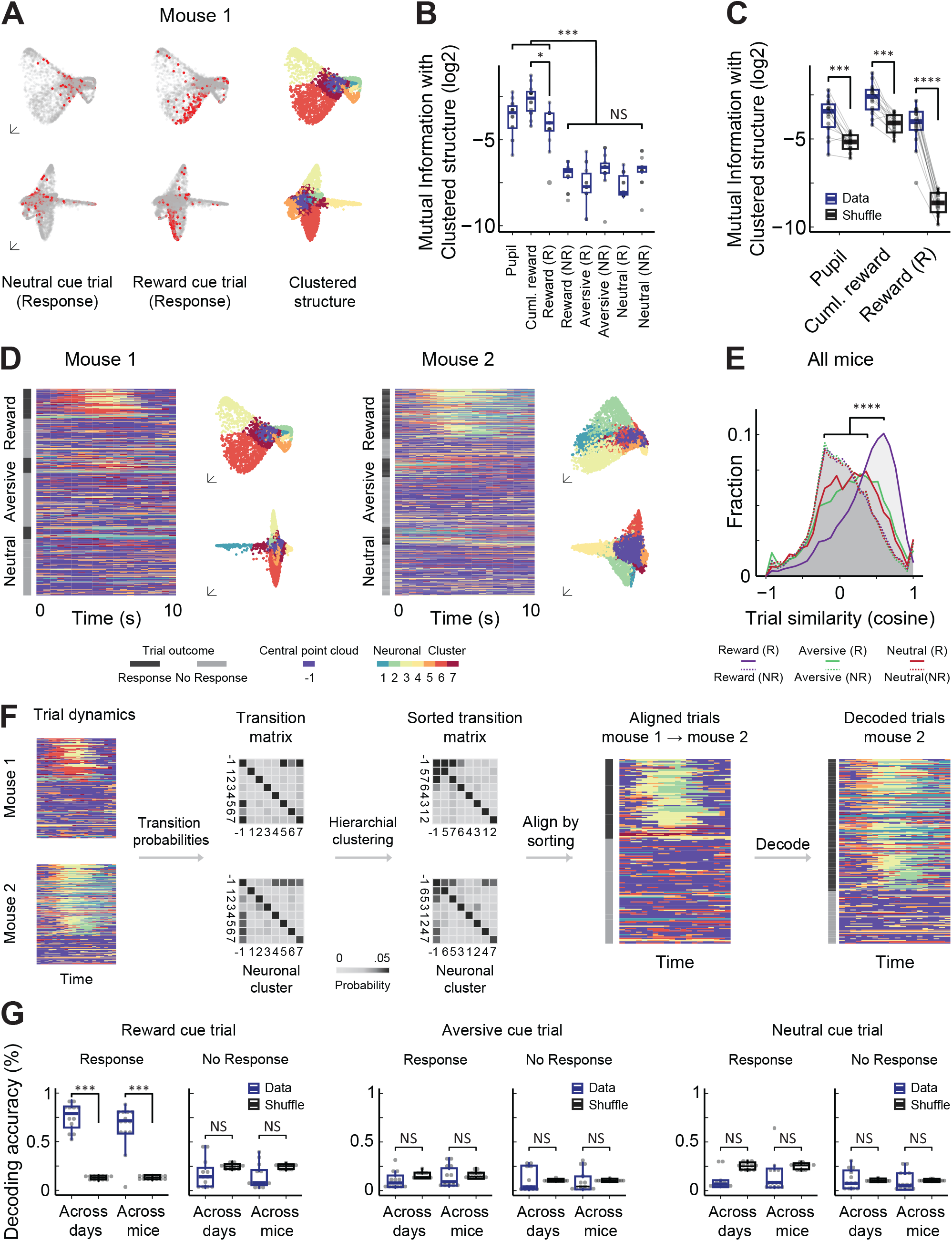
InsCtx activity dynamics within the manifold are stereotyped across different mice. **A.** Example visualization of time-points on the activity manifold that are associated with certain behavioral variables (red points). Different colors in the clustered structure reflect assignment to manifold clusters (**Figure S4**). Each row depicts a different plane of the activity manifold. **B.** Mutual Information between behavioral variables and the activity manifold. ***P<0.0004, NS: P<0.02, one-tailed Wilcoxon signed-rank test with Bonferroni correction. **C.** The most informative variables in ‘**B**’ exceeded chance levels, i.e., randomly shuffled labels (P<0.00095, one-tailed Wilcoxon signed-rank test). **D.** Example cluster sequence trial dynamics in two different mice. Left: two representative planes of the clustered structure (mouse 1 appears in ‘**A**’). Right: example of the cluster sequence dynamics for different trial types and outcomes (same cluster colors as the structures on the left). Note that for each mouse, cluster indices (colors) are randomly selected during the clustering procedure. **E.** Distribution of trial similarity (cosine similarity). ***P<5 10^−10^, two-sample Kolmogorov-Smirnov test with Bonferroni correction. **F.** Decoding procedure across datasets. Cluster labels are translated across datasets by aligning the order of the transition probability matrix between the different neuronal clusters. The translated dataset is then used for training to decode the different trial outcomes of the test dataset. **G.** Decoding accuracy for different trial types. ***P<0.00075 for across days and across mice versus shuffle, NS: P>0.2, one-and two-tailed Wilcoxon rank-sum test with Bonferroni correction, respectively. **A-G**. N=14 datasets from 5 mice

Using the clustered structure, we quantitatively assessed the relationship between the activity manifold and external/internal behavioral variables by computing their mutual information (**Figure S4D-E**; Methods). Three variables had higher mutual information with the activity manifold than others: (1) pupil size in inter-trial intervals (proxy for ongoing arousal levels^74^), (2) cumulative water rewards (proxy for slow changes in physiological state, i.e., water satiety^55^), and (3) trials in which the mice correctly responded to reward predicting cues to receive water rewards (referred to hereafter as “rewarded trials”; **Figure 2B**). Moreover, the mutual information for these three variables was significantly higher than that of shuffled data (**Figure 2C**). These results are consistent with previous work showing changes in InsCtx activity that are associated with rewards, arousal, and physiological state^20–22,50–52,55^. While we have previously shown that water rewards shift population activity along a linear thirsty-quenched axis^55^, manifold clusters associated with the quenched state were different from those associated with rewarded trials (**Figure S5A-B**). This suggests that there are also distinct representations of water satiety and reward in InsCtx population activity.

We next investigated activity manifold dynamics associated with behavioral variables. We represented each activity pattern by its manifold cluster, and then examined the temporal sequence of clusters during different trials (**Figure 2D**). InsCtx neuronal populations displayed somewhat stereotyped patterns of cluster transitions during rewarded trials, characterized by movement from the central point cloud to a sequence of transitions between the same 2-3 manifold clusters (**Videos S1, S2**). To evaluate the consistency of these dynamics, we calculated the pairwise similarity of cluster sequences across all trial types. Cluster sequences were highly similar across rewarded trials, substantially more so than all other trial types (**Figure 2E**). We confirmed these results using a different dimensionality reduction method (Isomap^75^; **Figure S6,S1F**). We also directly compared this cluster sequence method to our previous method of projecting activity onto linear thirsty-quenched axes^55^. As expected, the linear projection captures less of the variance of activity within the manifold and is less consistent across trials (**Figure S5C-D**).

Could this stereotyped sequence of activity patterns be used to decode trial outcomes across days and mice? The challenge is that our clustering technique does not yield functionally consistent cluster indices across mice. Thus, clusters associated with the same behavioral variable would not have the same label across mice. We therefore developed a method for translating cluster labels across dataset pairs (different days, different mice) to align activity dynamics between them for decoding (see Methods). We first determined the probability of transitions between different clusters in each dataset (**Figure 2F**, Transition matrix). We then translated the cluster labels from one “reference dataset” to relabel the clusters transition probabilities of the other dataset (**Figure 2F**, “Sorted transition matrix”). Finally, we used the translated cluster labels to train the decoder and tested it on the other “reference dataset” (**Figure 2F**). Using this approach, we could classify rewarded trials across different days (neurons were not aligned across days), and even across different mice, with high accuracy. We achieved an average accuracy of 75±13% and 65±13% (mean ± SD), respectively, which was ∼5 times higher than chance (∼12-15%). Decoding accuracy for other trial types did not significantly surpass that of shuffled trial labels (**Figure 2G**). We further validated these results using P-values derived from the shuffle distributions for each pairwise decoding (**Figure S7**). These results underscore that rewarded trials were consistently characterized by movement from the central point cloud to a sequence of transitions between the same 2-3 manifold clusters (**Videos S1, S2**).

Importantly, decoding trial types on single trials using this method was substantially more accurate than using average activity levels across all neurons, or using our previous approach of projecting activity on linear axes^55^ (**Figure S7**). Therefore, decoding accuracy for rewarded trials results from the stereotyped *patterns* of activity and does not trivially reflect global changes in activity levels.

### Insular cortex activity dynamics within the manifold are stereotyped across different motivational states and rewards

Given the remarkably stereotyped activity patterns we observed for rewarded trials in thirsty mice, and the consistent InsCtx manifold structure during both hunger and thirst, we next asked whether hungry mice would exhibit the similar dynamics with food rewards. We first visualized the temporal sequence of InsCtx manifold clusters in hungry mice and found that it displayed similar stereotyped sequences as in thirsty mice (**Figure 3A**). In contrast, activity sequences in visual areas were very different, primarily tracking visual cues, independent of rewards (see also **Figure S3**). We then tested whether we could decode trial outcome across different thirsty and hungry mice, receiving water and food rewards, respectively. Decoding accuracy in *hungry* mice was high when training the decoder on InsCtx activity from *thirsty* mice (67±23%, mean ± SD, vs. ∼12-15% chance). However, decoding accuracy was near chance when training the decoder on activity from visual areas (11±12%; **Figure 3B**). Notably, this differential decoding was not due to the clustering procedure as InsCtx and visual areas datasets both had 7 clusters. The consistent activity dynamics in InsCtx across hunger and thirst suggest that these activity patterns likely do not reflect salient sensory attributes of the reward that are different in water vs. Ensure (e.g., taste, viscosity), or the underlying physiological need. Nevertheless, we cannot rule out that other sensory features, common to Ensure and water rewards, could be represented in these activity patterns.

**Figure 3–.**
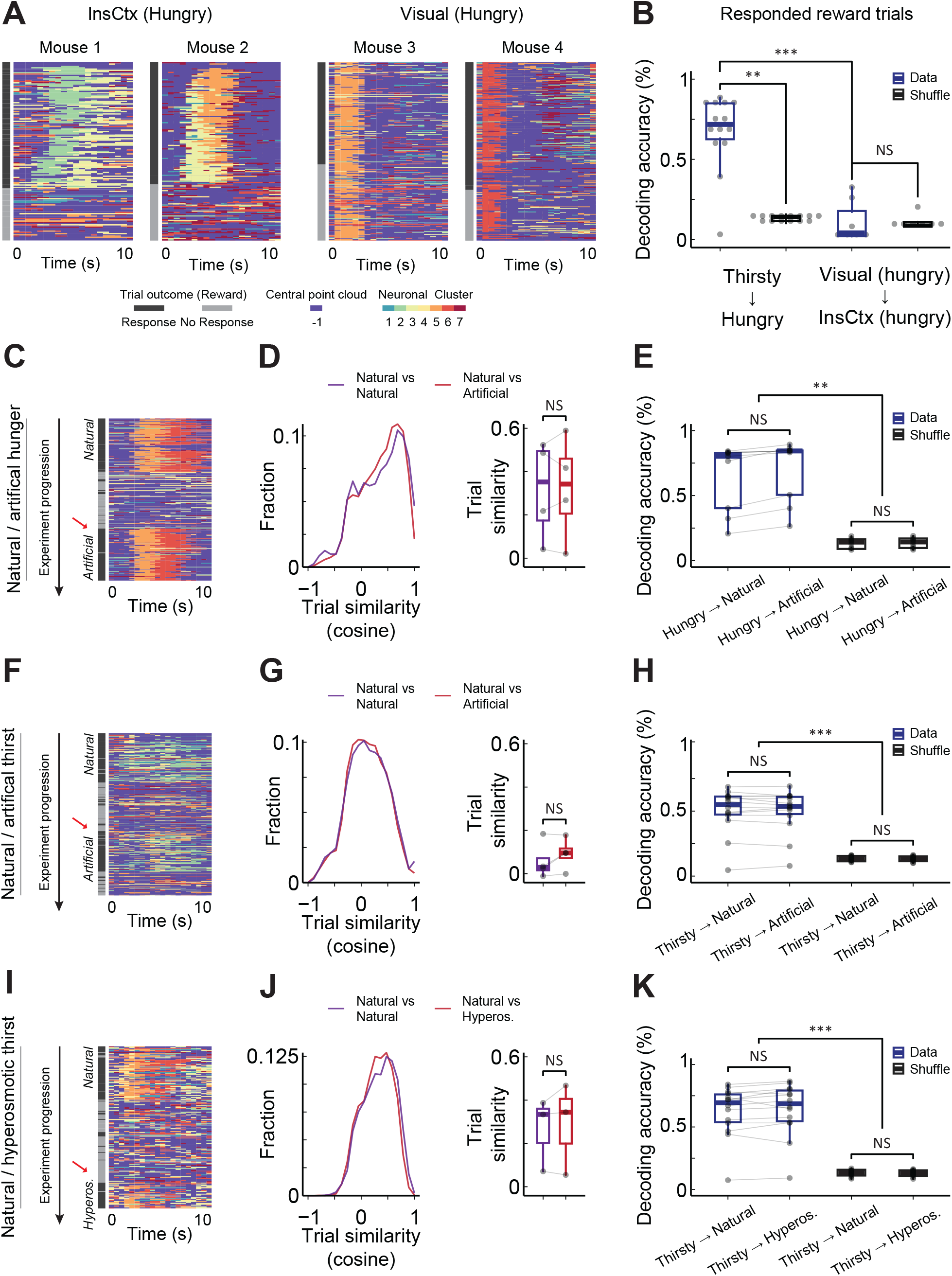
InsCtx activity dynamics within the manifold are stereotyped across different motivational states and rewards. **A.** Visualization of reward cue trial dynamics in two hungry mice from InsCtx and visual areas. **B.** Decoding accuracy of rewarded trials from hungry mice. The decoder was trained using datasets from InsCtx in thirsty mice or from visual areas in hungry mice. ***P<0.0002, **P<0.0065, NS: P>0.35. Dunn test with Bonferroni correction. N=9 datasets from 6 mice. **C.** Left: Example activity dynamics of reward cue trials before and after artificial induction of hunger using chemogenetic activation of AgRP “hunger neurons”. Red arrow: first responded reward cue trial following induction. **D.** Distribution of pairwise trial similarity (cosine similarity) among rewarded trials across natural and artificially induced hunger. Left: distributions pooled from all datasets. Right: paired comparisons of the mean pairwise trial similarity for each dataset trials. NS: P>0.15, two-tailed paired t-test. **E.** Comparison of the decoding accuracy of rewarded trials across natural and artificially induced hunger. Comparisons between natural and artificial data across all states: P>0.3. Comparisons between natural and artificial shuffles across all states: P>0.08. **P<0.008. One-tailed Wilcoxon signed-rank test with Bonferroni correction. **F-H**. Same as in ‘**C**’-‘**E**’ before and after artificial induction of thirst using chemogenetic activation of SFO “thirst neurons”. **I-K**. Same as in ‘**C**’-‘**E**’ with before and after induction of hypertonic thirst using injection of hypertonic saline. **A-K**. N=10 hungry datasets from 7 mice, N=14 thirsty datasets from 5 mice, N=7 visual datasets from 5 mice, N=4 AgRP activation datasets from 4 mice, N=4 SFO activation datasets from 4 mice, N=3 hypertonic saline injection datasets from 3 mice.

We further tested the extent to which these activity patterns remain stereotyped in mice performing similar behavioral tasks across additional natural and artificial motivational states. Specifically, we examined InsCtx activity from mice in which artificial hunger or thirst were induced by chemogenetic activation of hypothalamic AgRP “hunger neurons” or SFO^GLUT^ “thirst neurons” − the primary sensors of physiological deficits and actuators of relevant behavioral and physiological responses^76–79^. We also analyzed InsCtx activity from mice in which we induced hyper-osmotic thirst by administration of hypertonic saline^80^. These datasets gave us the unique opportunity to compare, within each mouse, the activity during the natural motivation driven by physiological need, with the artificially induced motivation.

We first visualized the sequences of manifold clusters and compared them during the natural motivational state to after induction of the artificial motivational state. Sequence transitions during rewarded trials were remarkably similar in both states (**Figure 3C,F,I**). We quantified this using pairwise trial similarity before and after induction of the artificial motivation. For all motivational states, the pairwise similarities remained similar between the natural and artificially-induced states, with no significant changes (**Figure 3D,G,J**). We therefore trained a decoder on datasets from hungry and thirsty mice, which were not part of the artificial motivation experiments, and used these to test decoding before and after the artificial induction of motivational states. Decoding accuracy for rewarded trials remained the same for both states, and was significantly higher than chance (**Figure 3E,H,K**). These findings demonstrate that the stereotyped activity patterns in InsCtx are not specific to a particular reward (including its sensory properties), nor to the specific underlying motivational state. Importantly, closer examination of activity patterns and of individual neurons, will likely reveal distinct representations of different rewards and motivational states. Nevertheless, our analyses reveal a common activity pattern, which is so robust as to enable single-trial decoding across individuals and motivational states.

### Insular cortex stereotyped activity dynamics reflect goal-directed reward anticipation, but not licking, taste, or positive valence

The stereotyped activity patterns we observed in InsCtx across rewarded trials with different rewards and motivations could reflect goal-directed behavior, licking, positive valence, or need fulfilment. We next tested these different options.

In hungry and thirsty mice, we observed that the stereotyped cluster sequence during rewarded trials appeared linked with anticipatory licking, as evidenced by sorting trials based on the onset of anticipatory licking (**Figure 4A**). We thus quantified the correlation between the onsets of sequential activity and anticipatory licking in individual trials (**Figure 4B**). These correlations were notably high and statistically significant in hungry mice (0.5±0.13; mean ± SD) and thirsty mice (0.47±0.15; **Figure 4C**). Additionally, the onset of anticipatory licking consistently preceded the onset of sequential cluster activity, and importantly, occurred independently of reward delivery (**Figure 4D-E**). Together with similar activity dynamics across food and water rewards (**Figure 3**), this supports a dissociation from the sensory aspect of the reward. Thus, these stereotyped patterns could potentially reflect licking per se, positive valence (independent of the reward’s sensory properties), or cue-driven goal-directed behavior.

**Figure 4–.**
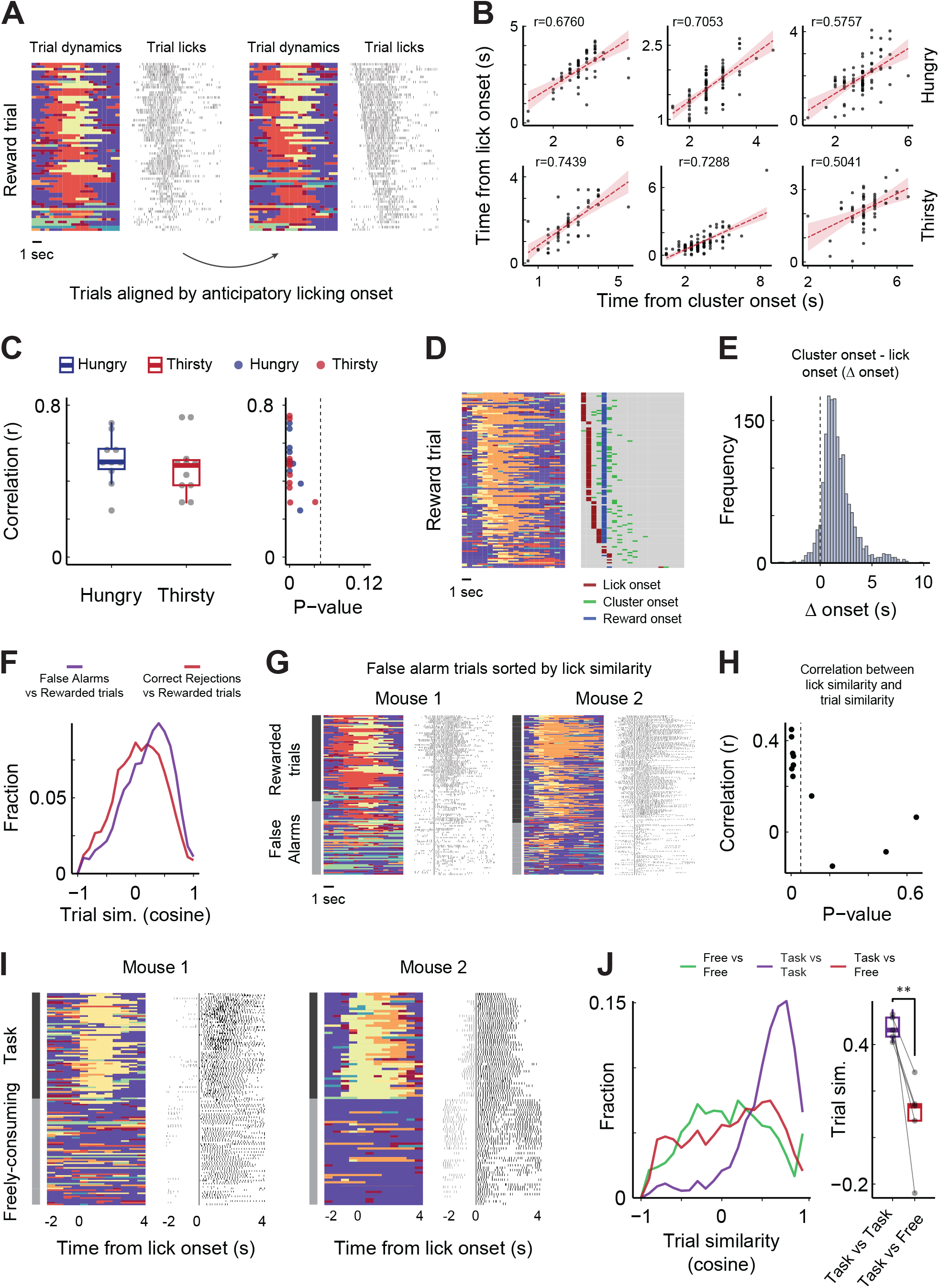
Stereotyped activity dynamics reflect goal-directed reward anticipation, but not licking, taste, or valence. **A.** Example neuronal dynamics during rewarded trials and task-related licking behavior. **B.** Example correlations between anticipatory licking onsets and rewarded trials cluster onsets in 6 datasets from hungry (top) and thirsty mice (bottom). **C.** Left: Distribution of all correlations across all hungry (N=10) and thirsty (N=11) datasets. Right: Scatterplot of correlation levels vs. p-values. Dashed vertical line: P=0.05. P<0.04, for all datasets, Pearson’s product-moment correlation. **D.** Example visualization of reward onset, anticipatory licking onset, and activity cluster onset. Left: Cluster dynamics of all rewarded trials. Right: Onsets of anticipatory licking, reward delivery, and cluster dynamics per trial. Note that the onset of the sequential cluster pattern consistently follows the onset of anticipatory licking and appears unrelated to the onset of reward delivery. **E.** Distribution of the time difference between rewarded trial cluster onset and anticipatory licking onset. Across all trials, cluster onset occurs approximately 1 second *after* lick onset. **F.** Distribution of trial similarity between incorrectly responded aversive and neutral trials (“false-alarms”) and rewarded trials (purple), compared to the similarity between correctly non-responded aversive and neutral trials (“correct-rejections”) to rewarded trials (red). N=14 datasets from 5 thirsty mice. **G.** Two examples of cluster sequence trial dynamics during false-alarm trials and rewarded trials. False alarm trials were sorted based on their *licking* pattern similarity to the *licking* patterns in rewarded trials. Note that as licking pattern becomes more similar, false-alarm cluster sequence trials become more similar to rewarded trials. **H.** Distribution of all correlations between licking similarity and trial similarity in false-alarm trials and rewarded trials, as a function of p-value. 9/11 had positive correlations, 7/9 were statistically significant (P<0.05, Pearson’s product-moment correlation). **I.** Example cluster sequence dynamics and licking behavior in two mice during reward consumption in the operant task and during self-paced free consumption. Dynamics were aligned based on the onset of licking. **J.** Left: Distribution of all pairwise comparisons between reward consumption trials during task engagement and free consumption behavior (N=5 datasets from 5 mice). Right: Comparison of average trial similarity for each dataset individually. **P<0.009, one-tailed paired t-test.

To distinguish between these possibilities, we examined trials in which mice incorrectly licked to cues that do not predict reward (“false-alarm” trials). Cluster sequences in false-alarm trials resembled rewarded trials more than other trial types (**Figure 4F**). To further investigate this, we directly compared cluster sequences for false-alarm and rewarded trials. We sorted false-alarm trials based on the similarity of *licking* patterns to rewarded trial licking (**Figure 4G**). False-alarm trials with similar licking patterns to rewarded trials also displayed similar cluster sequence dynamics. We quantified this by assessing correlations between licking similarity and cluster sequence similarity in false-alarms vs. rewarded trials, finding positive correlations in 9/11 datasets (**Figure 4H**). As mice are not actually rewarded in false-alarm trials, these results support the conclusion that InsCtx stereotyped activity patterns do not reflect reward or positive valence of cues or outcomes. Nevertheless, they could still trivially reflect licking, or cue-associated reward expectation.

To differentiate between these possibilities, we compared cluster sequences in hungry mice during reward consumption in the operant discrimination task vs. during free consumption in the same context without predictive cues or task structure (head-fixed, visual cue monitor on without cues). Notably, in both conditions mice had a similar physiological need, fulfilled using the same action (licking), to obtain the same reward. We constructed ‘pseudo-trials’ during the free consumption epochs by identifying significant breaks between lick bouts and analyzed the corresponding sequences of manifold clusters. As expected, average population activity increased during reward consumption in both task-structured trials and self-paced consumption (**Figure S8**). However, activity dynamics during task engagement and free reward consumption were distinctly different (**Figure 4I**). Quantitatively, pairwise trial similarity between operant trials and free consumption trials was significantly lower than the pairwise similarity within operant trials. (**Figure 4J, S8**). We verified that these results could not be explained by differences in licking patterns (**Figure S8**). Specifically, we compared licking patterns across task performance and free consumption, that are as similar as possible (and statistically indistinguishable). We still found the same dramatic differences in activity dynamics (**Figure S8**). However, we cannot rule out the existence of subtle differences in licking patterns between these two conditions. It seems unlikely that such subtle differences, rather than behavioral context, account for such dramatic differences in InsCtx population activity patterns. Future work could directly test this.

In summary, by comparing highly similar licking patterns across different behavioral contexts (rewarded trials, false-alarm trials, free reward consumption), we show that: (1) licking per se cannot explain the stereotyped activity patterns we observed (comparing rewarded trials vs. free reward consumption); and that (2) reward expectation regardless of reward receipt does explain the stereotyped activity patterns we observed (comparing rewarded trials vs. non-rewarded false-alarm trials). Taken together, these results suggest that the stereotyped sequential activity patterns we observed in InsCtx are closely related to learned anticipation of reward during goal-directed behavior, rather than licking, the sensory aspects of the reward, positive valence, or specific motivational states.

## Discussion

We used unbiased unsupervised machine learning and topological approaches to discover that InsCtx population activity had consistent dimensionality, manifold structure, and dynamics across days, across different mice, and across different motivational states.

Most cellular level InsCtx neural recordings in behaving animals have focused on taste processing^26,43,44,46^. These studies revealed sequential temporal coding of tactile, chemosensory, and palatability information in individual neurons on the order of hundreds of milliseconds^26^. Small populations of 5-10 InsCtx neurons have been shown to exhibit metastable dynamics, transitioning between different states that can reflect different behavioral variables, each lasting tens to hundreds of milliseconds^33–35^. Other work using activity manipulations and bulk activity recordings has shown that InsCtx activity is related to taste, palatability, motivation, valence, and anxiety^20,22,32,36,52,53,81^. Future experiments using manifold analyses of large population activity should be designed specifically to capture these variables, whether on slow or fast time-scales. The faster stereotyped changes we discovered appear not to be related to licking, taste, positive valence, or general reward. These observations (**Figure 4**) may explain previous manipulation experiments, in which inhibiting InsCtx activity reduces task-related behaviors, while not affecting free consumption behavior ^50,82^. Future work could test this interpretation.

A recent study found that activity of anterior InsCtx (deep layer) projections to the brainstem encodes motivational vigor^52^. Notably, the stereotyped activity patterns we described here in superficial layers 2/3 of mid InsCtx likely do not reflect motivational vigor per se, as they depended on behavioral task structure and did not change with partial satiation, which reduces motivational vigor^52^. Nevertheless, it will be important to understand how InsCtx can simultaneously represent different types of information, and to what extent they are eventually integrated before being relayed to downstream brain regions to affect behavior and bodily physiology. Indeed, different InsCtx projection populations (e.g., projecting to striatum, thalamus, or brainstem^8,61,62^), different InsCtx layers (e.g., superficial vs. deep), and different InsCtx subregions (e.g., granular/dysgranular/agranular, and anterior/mid/posterior), may convey different types of information to downstream regions.

We previously used linear projections in activity space to suggest that reward consumption transiently shifts activity towards a future satiety state^55^. Here, using analyses of the entire activity manifold we find stereotyped goal-directed reward anticipation activity patterns, which are mostly independent of changes in ongoing activity that reflect physiological state. Thus, while our previous hypothesis-driven work provided support to a dominant model of interoception^5,7^, our unbiased analyses here highlight a general computation InsCtx performs, which could be important for many behaviors that do not directly involve interoception.

Activity dynamics on the manifold were remarkably consistent across individuals and motivations, suggesting that these dynamics may reflect an important fundamental aspect of InsCtx function. We speculate that this is related to the consistent involvement of InsCtx in cravings during pathological conditions associated with aberrant reward processing, including obesity, binge eating disorder, and drug addiction^1–4,8^. InsCtx has been suggested to encode the anticipated interoceptive value, which when goes awry can lead to aberrant food or drug cravings^1^. We thus speculate that the stereotyped goal-directed reward anticipation patterns we discovered reflect this fundamental computation in InsCtx. As such, future work using the same analytical framework in mouse models of pathological conditions, could test this idea to form a deeper understanding of the role of InsCtx in pathological conditions that involve aberrant motivations.

### Limitations of the study

We developed and used one manifold analyses approach. Future studies should compare our methods to other recent powerful methods^12^. Additionally, our analyses were limited by the behavioral variables that were measured in our studies, and the specific behavioral context of cue-driven reward consumption. Other parts of the manifold could be determined by the intrinsic connectivity of the network, as has been previously suggested^35^, and/or by other variables we did not measure in these experiments^21,42^. Furthermore, it will be important to compare InsCtx activity across different behavioral contexts, including those that do not involve reward consumption, to assess its intrinsic dimensionality and structure of the manifold.

## Acknowledgements

We thank Mark Andermann, Alon Rubin, Yarden Cohen, Sasha Devore, Stav Shtiglitz, Ayal Lavi, Yael Prilutski, Einav Litvak, Omer Izhaki, Inbar Perets, Daniel Deitch, Tom Talpir, Omer Richmond, Gal Elyasaf, Birgit Jickeli, and Daphna Nachmani for fruitful discussions and helpful comments on the manuscript. YL is supported by research grants from the Israel Science Foundation (ISF #860/21), the European Research Council (ERC StG #101039145), and the Center for New Scientists at the Weizmann Institute of Science.

## Author contributions

IT and YL conceived, designed, and executed the study. IT analyzed the data. IT and YL wrote the manuscript.

## STAR Methods

### Key resources table

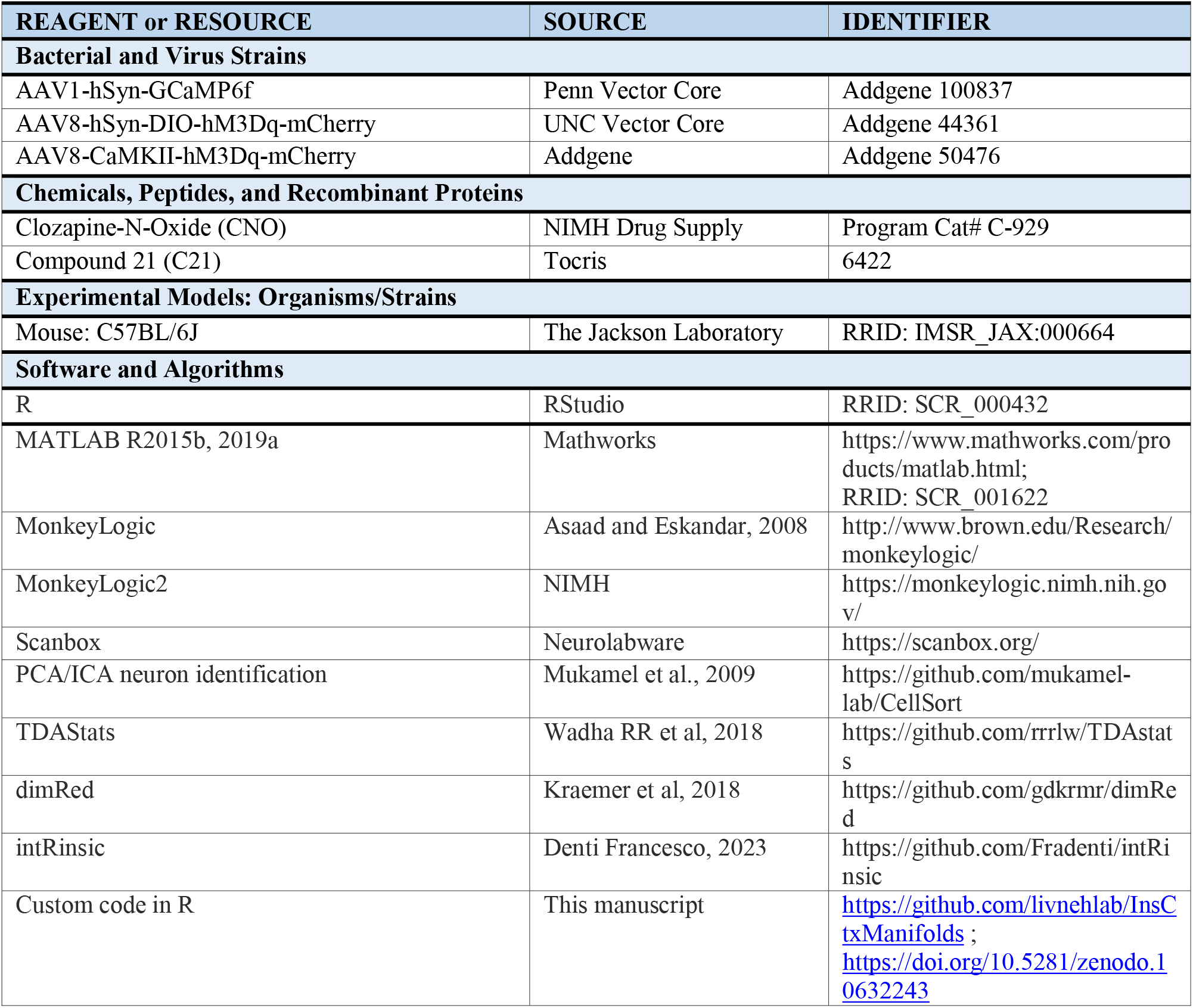

### Resource availability

#### Lead contact

Further information and requests for resources and reagents should be directed to the lead contact, Yoav Livneh.

### Material availability

This study did not generate new unique reagents.

### Data and code availability

The code used for the analyses in this work is available at the LivnehLab github (https://github.com/livnehlab/InsCtxManifolds; https://doi.org/10.5281/zenodo.10632243).

Any additional information required to reanalyze the data reported in this paper is available from the lead contact upon request.

All data reported in this paper will be shared by the lead contact upon request.

### Experimental model and study participant details

All animal care and experimental procedures were approved by the relevant Institutional Animal Care and Use Committee. Mice used for *in vivo* two-photon imaging (males, age at surgery: 9-15 weeks) were instrumented with a headpost and a 2 mm microprism, centered over the mid InsCtx (see details below).

### Method details

Datasets analyzed for this paper were both previously published data^50,55,73^, as well as unpublished data. Unpublished datasets included experiments similar to those described in the published datasets (e.g., in water or food restricted mice performing an operant visual discrimination task for water or Ensure rewards), as well as imaging during head-fixed satiation by Ensure consumption and induction of hyperosmotic thirst by intraperitoneal injection of hypertonic saline. We briefly describe the experiments below, see also refs^50,55^ for more detailed descriptions.

### Behavioral task

To perform the behavioral task, mice were water restricted to ∼80% of their pre-restriction body weight or food-restricted to ∼85% of their free-feeding body weight. We trained mice to discriminate between square-wave drifting gratings differing in orientation presented on an LCD screen (2 Hz and 0.04 cycles/degree, full-field square wave drifting gratings, 80% contrast; food cue: 0°, aversive cue: 270°, neutral cue: 135°^50,55^. All drifting gratings were presented for 2 sec, after which the mouse had a 2 sec window to respond with a lick. Licking during the visual cue was not punished, but also did not trigger delivery of the water/Ensure/salt-water/quinine. Only the first lick (if any) occurring during the response window triggered delivery of water/Ensure/salt-water/quinine. The lickspout was designed with two adjacent lick tubes (one for each outcome), such that the tongue contacted both tubes on each lick, which served as an effective deterrent for lick responses following aversive cues. Well-trained mice had a high rate of correct water/food cue licking responses (criterion: >80% of trials, usually ∼90-95%), and a low rate of licking following aversive cue presentations (criterion: <50%, usually ∼20-30%). Each water reward was a ∼2-3 µL drop, and Ensure reward was a ∼5 µL drop (0.0075 calories). Behavioral training was performed using MonkeyLogic^83^ and MonkeyLogic2 (https://monkeylogic.nimh.nih.gov/).

### Surgical procedures

Stereotaxic injections AAV8-CaMKII-hM3Dq-mCherry, AAV8-hSyn-DIO-hM3Dq-mCherry, and AAV1-hSyn-GCaMP6f, as well as and implantation of microprisms (2 mm prisms; #MCPH-1.0; Tower Optical; coated with aluminum along their hypotenuse) were performed as previously described^50,55^.

### Two-photon imaging across different natural and artificial states

Two-photon imaging of GCaMP6f was performed using a resonant-scanning two-photon microscope with tiltable scanhead (Neurolabware; 31 frames/second; 1154×512 pixels). All imaging was performed with a 20x 0.45 NA air objective (Olympus) with a 540 x 360 μm2 field of view. All imaged fields of view (FOV) were at a depth of 90-200 µm below the pial surface, using a Mai Tai DeepSee laser (Newport Corp.) with laser power at 920-960 nm of 35-80 mW at the front aperture of the objective (power at the sample was likely substantially less due to partial transmission via the microprism). Imaging depth was adjusted in between runs (every 30 min) to account for slow drift in the z plane (< 7 μm). Recording locations were approximately +0.5 mm to −1.0 mm to the middle cerebral artery (see^50,55^ for further details).

### Imaging across thirsty and quenched states

We imaged mice during gradual water satiation during consecutive 30-minute runs until the mice voluntarily stopped performing the task. We then performed one more imaging run, the ‘quenched’ run, in which mice did not respond to the water cue.

### Imaging across hungry, satiation, and sated states

We imaged mice in two blocks of trials within a session, one block during food restriction and a subsequent block immediately following re-feeding. At the start of each imaging session, food-restricted mice performed the visual cue discrimination task. After ∼180 trials (30-min imaging run), we provided the mouse with ad libitum access to Ensure until voluntary cessation of consumption. Ensure consumption lasted 45-75 minutes. We triggered delivery of Ensure with every lick, but with a minimum inter-trial interval of 2.5 s between Ensure deliveries. During this period of time, mice consumed ∼3-5 mL of Ensure and then voluntarily stopped licking for rewards. We then imaged additional ∼180 trials (30 min imaging run) while mice were satiated (operationally defined as the absence of voluntary licking).

### Imaging during chemogenetic activation of SFO^GLUT^ neurons

Following imaging during the quenched state (see above), we injected 150 µl 0.9% of either saline or CNO (5 mg/kg), waited 10-15 min and started another imaging run (∼180 trials, 30 min). For every mouse used for these experiments, we used postmortem histology and immunohistochemistry to verify hM3Dq-mCherry expression in the SFO.

### Imaging during following injection of hypertonic saline to induce hyperosmotic thirst

Following imaging during the quenched state, we injected ∼200 µL of 2M NaCl, waited 5-10min and started another imaging run (∼180 trials, 30 min). Mice usually re-engaged in the behavioral task immediately or after 2-3 minutes.

### Imaging during chemogenetic activation of AgRP neurons

Following imaging during hunger and satiety, mice were returned to their home-cage with ad libitum access to regular chow. The next morning, we imaged the same InsCtx field of view in this satiety state (100-110% of normal body weight) during ∼180 trials (30 min.). We then injected CNO (1-3 mg/kg). Ten minutes later, we initiated an additional imaging run of ∼180 trials. For every mouse used for these experiments, postmortem histology and immunohistochemistry confirmed hM3Dq-mCherry expression in the hypothalamic arcuate nucleus.

### Pupil videography during two-photon imaging

We acquired data using a GigE Vision camera (Dalsa) with a 60 mm lens (Nikon MicroNikkor) from a pre-selected region of interest around the eye ipsilateral to the LCD monitor used to present visual cues (contralateral to the InsCtx microprism). Acquisition of each frame (frame rate of 15.5 Hz) was triggered on every other frame of two-photon acquisition (acquired at 31 Hz) using Scanbox software (Neurolabware). The pupil was backlit with illumination originating from diffusion within the brain of the IR light used for two-photon excitation during imaging. See below for details of data analysis.

### Image registration and time course extraction

First, each acquired image was spatially down sampled by 2X. To correct for motion along the imaged plane (x-y motion), each frame was registered to an average field-of-view using efficient subpixel registration methods^84^ Within each imaging session, each run (2-8 runs/session) was registered to the first run of the day. Image stacks were de-noised using principal components analyses (PCA) of every pixel across time, and by user identification and removal of noise principal components (low eigenvalues; based on^85^). Cell masks and calcium activity time courses (‘F(t)’) were extracted using custom implementation of common methods^85^. To avoid use of cell masks with overlapping pixels, we only included the top 75% of pixel weights for a given mask, but users screened each prospective ROI and could edit the size of the mask, selectively removing the lowest probability pixels. We then excluded any remaining pixels identified in multiple masks. We manually verified that all cell masks had typical cell body morphology and size.

Fluorescence time courses were extracted by averaging the pixels within each region-of-interest (‘ROI’) mask. Fluorescence time courses for neuropil within a 25 µm annulus surrounding each ROI (but excluding adjacent ROIs and a protected ring surrounding each ROI) were also extracted (F_neuropil_(t): median value from the neuropil ring on each frame). Fluorescence timecourses were calculated as F_neuropil_corrected_(t) = F_ROI_(t) - F_neuropil_(t). The change in fluorescence was calculated by subtracting a running estimate of baseline fluorescence (F_0_(t)) from F_neuropil_corrected_(t), then dividing by F_0_(t): ΔF/F(t) = (F_neuropil_corrected_(t) - F_0_(t))/ F_0_(t). F_0_(t) was estimated as the 10th percentile of a 32 sec sliding window^50,55,73^.

### Preprocessing of neuronal data

For each collected dataset, we down-sampled ΔF/F traces by averaging every consecutive 15 frames (acquired at 31 Hz), resulting in ∼0.5 seconds time-bins. Subsequently, we excluded outlier neurons with abnormally high ΔF/F values, calibrated separately for two different microscopes (2000% and 5000%). Following this, we Z-scored the activity time course for each individual neuron. Each dataset consisted of multiple 30-minute runs, acquired within the same day. For certain datasets, we applied separate Z-scoring to each run if substantial extended breaks occurred between runs or if there was a z-plane shift between the runs.

### Intrinsic dimensionality estimation

To estimate the intrinsic dimensionality of the neuronal activity data, we employed the minimal neighborhood information technique as an estimator^86^. Briefly, the intrinsic dimension is estimated using the ratio formula:

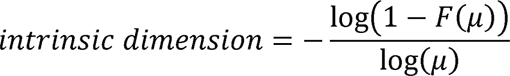

Here, *μ_i_* was determined for each data point *i*, where 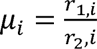, with *r*_1,*i*_ and *r*_2,*i*_ representing the Euclidean distances of the nearest and second-nearest data points to point *i*, respectively. *F*(*µ_i_*) indicates the percentile of *μ_i_* with respect to all other data points. Subsequently, *μ_i_* was computed for each data point, and the highest 4% of points were excluded. The intrinsic dimension was defined as the slope calculated from a linear regression between log(1-*F*(*µ*)) and log(*µ)* across all data points. To better estimate the intrinsic dimensionality of the neuronal activity data, we used dimensionality reduction for denoising prior to dimensionality estimation. In essence, dimensionality estimation remains invariant to the embedding dimension. To ensure this, we iteratively assessed the intrinsic dimension across intermediate dimensionality reductions (see **Figure S1B-E**). We further verified this in certain datasets where the number of neurons was lower (specifically in visual areas from hungry mice; see **Table S1**). In visual areas datasets, we compared the estimated dimensionality with InsCtx hungry-sated datasets (see **Figure 1**). To ensure that dimensionality was unaffected by the number of neurons, we repeatedly randomly subsampled to the lowest number of neurons (see **Table S1**). This confirmed that the estimated dimensionality was consistently lower in InsCtx vs. visual areas. Specifically, for visual areas, the average estimated dimension was 11.7±3 (mean ± SD) for both subsampled and full data. In contrast, hungry-sated datasets, this resulted in 7.7±2 and 8.7±2 for subsampled and full data, respectively.

### Non-linear dimensionality reduction

To reduce the dimensionality of the neuronal data matrix *X*, we applied Laplacian eigenmaps^64^ implemented by a pre-established pipeline^87^. Laplacian eigenmaps, a spectral non-linear dimensionality reduction technique, operates on a data matrix *X* with dimensions *N* x *T*, by generating a weighted adjacency graph *W* for the *T* high-dimensional data points such as each time point *t* ∈ ℝ*^N^*. This adjacency graph is computed for all pairs of data points, where the weight between a pair of data points *x_i_*, *x_j_*, ∈ *X* is assigned as *W_ij_* = 1 if the data points are connected and *W_ij_* = 0 otherwise. Data points are considered connected if they fall within the *K* nearest neighbors of each other, where *K* is a user-defined parameter (see below). Following the computation of *W*, a diagonal weight matrix *D* is constructed with *D_ii_* = ∑*_j_ W_ji_*, which is equivalent to aggregating the rows of *W*. Notably, the Laplacian of the original data matrix is *L* = *D* - *W*. The process of obtaining the low-dimensional embedding is achieved by calculating the eigenvalues and eigenvectors of *L**f*** = *λD**f***. After excluding the leading eigenvector^64^, the following *m* eigenvectors are utilized to obtain a new low-dimensional mapping for each data point *x_i_*, which is given by^17^

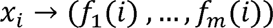

with *m* representing the desired output dimensionality. For every dataset *X*, we conducted two rounds^41^ of dimensionality reduction using the algorithm described above. In the initial iteration, a 20-dimensional matrix was generated, with a parameter *K* set to 7.5% of the time points in matrix *X*. The second iteration was performed on the 20-dimensional matrix obtained from the preceding step. This culminated in the matrix reaching its ultimate dimension – the mean estimated intrinsic dimension of 6 – with a parameter *K* set to 2.5% of the time points in matrix *X*. The mean estimated intrinsic dimension of 6 was derived from the process described above.

### Topological features analysis and topological similarity

To assess the topological features of the data, we quantified for each dataset its components (β0) and holes (β1) as a function of the radius threshold, using a pre-established algorithm^67^ (TDAStats: R pipeline for computing persistent homology in topological data analysis, https://github.com/rrrlw/TDAstats). Due to computational considerations, we determined the topological characteristics of clustered timepoints within the reduced data matrix, acquired through K-means clustering with a chosen *K* value of 80. Given our utilization of a stochastic variant of K-means, this process was repeated 20 times for each dataset. For every topological feature extracted from the clustered data matrix, we quantified its lifespan and derived the lifespan distribution for all topological features. The topological resemblance between a pair of datasets was defined as the mean Wasserstein distance^88^ between the lifespan distribution of features from both datasets, covering all 20 iterations of K-means clustering. To ascertain the significance of differences between dataset pairs, permutation tests were conducted, following a methodology outlined in prior work^69^. The lifespan distributions of a pair of datasets were pooled and then systematically rearranged to generate two separate subsamples, adhering to the original sizes of each distribution. The Wasserstein distance was computed between these subsamples. This procedure was repeated n=200 times, and a P-value was established as the percentage of permutations yielding a Wasserstein distance less than the original distance between the dataset pairs. The ultimate P-value for every pair of datasets was determined as the average of the P-values derived from this process across all 20 K-means clustering iterations.

### Clustering the activity manifold

To evaluate the activity manifold, our objective was to create a standardized comparison across datasets through an automated clustering approach. To achieve this, we devised an unsupervised clustering strategy that recognizes data points that deviate from the central point cloud, and thus classifies them as distinct neuronal clusters. To do so we systematically excluded time points that were closest to the centroid of the reduced data matrix. Starting from 40 times points and progressing to the entire dimensionality of the data matrix. For each set of excluded timepoints, we conducted 20 iterations of K-means clustering with *K* values spanning from 2 to 20. The mean squared error (MSE) was computed between the resulting clusters and their centroids across all iterations and for each *K* value. This procedure produced a two-dimensional grid with its two dimensions being the fraction of excluded timepoints and the chosen *K* values for K-means clustering. The cells of the grid contained MSE values over the 20 clustering repetitions. To identify the optimal configuration, we scaled both grid dimensions to 1 and determined the grid cell closest to the origin of a three-dimensional Cartesian system formed by the grid, where the third dimension was the MSE value within each cell. This process yielded, for each dataset an optimal configuration consisting of 2 parameters: a *K* value, and the number of datapoints to be considered as the central point cloud. Across virtually all utilized datasets the identified optimal *K* value was 8, except one dataset in which it was 7. After obtaining the optimal configuration, our aim was to derive deterministic clustering labels for each dataset. This was accomplished by implementing the optimal configuration, isolating timepoints deemed part of the central point cloud, and executing 500 iterations of K-means clustering with the optimal K value for the remaining timepoints. The resulting clustering labels were aggregated into a 500 x *T* matrix. This matrix was then subjected to an additional clustering step using the ward agglomerative complete linkage hierarchical clustering method^89^. This yielded the final labels of the clustered activity manifold.

### Activity manifold parameterization

To evaluate the contribution of behavioral variables to the activity manifold, we devised labeling for the distinct behavioral variables, considering the total timepoints under analysis. These variables were categorized into two groups: discrete (binary) and continuous. Discrete binary variables encompassed the potential classes of trial outcomes within the behavioral task (e.g., response vs. no response for a given trial type). For each of the six trial outcome classes, a corresponding label vector was generated, spanning *T* timepoints. A value of *Trail outcome Label* = 1 was assigned for timepoints ranging from 3 to 6 seconds after the cue onset for each class, while a value of *Trail outcome Label* = 0 was allocated to the remaining timepoints. This enabled the isolation of timepoints exclusively linked to that specific trial outcome, excluding those related to visual cues. As for continuous variables, namely Pupil Size and Cumulative Consumed Rewards, their magnitude across the entire experiment was quantified. Pupil size was quantified using code from previously published work^90^. Two label vectors, each with a length of *T*, were established for these variables. To ensure equitable comparison with discrete variables, both Pupil Size and Cumulative Consumed Rewards were divided into eight bins.

### Mutual information between behavioral variables and the activity manifold

To determine the mutual information between behavioral variables and the activity manifold, we constructed a joint contingency table that related the two label vectors: *Cluster* and *Behavior*. Here, *Cluster* represents a vector of size *T* containing different neuronal cluster labels (including the central point cloud), while Behavior was a *T*-sized vector representing behavioral variables (ranging from 0 to 1 for discrete variables and 1 to 8 for continuous variables). Following this, we utilized the contingency table to formulate a joint probability space, denoted as *P_Cluster_*_x*Behavior*_. The calculation of mutual information was carried out according to the formula:

Where *P_Cluster_* and *P_Behavioor_* are the respective marginal probability distributions. To reduce potential variations caused by different behavioral factors and ensure an equitable comparison with discrete variables, for the continuous variables we exclusively included time points unaffected by behavior, following established procedures from our previous work^55^. These time points corresponded to the final 3 seconds of inter-trial intervals following the correct rejection of aversive and neutral trials (referred to as non-responded trials).

### Trial structure manifold dynamics

To evaluate manifold dynamics in relation to the trial structure, we compiled the sequential neuronal cluster label (including the central point cloud) time course for every trial. This was achieved by capturing a 10-second interval from the onset of the cue for each trial class. They yielded a sequence of 20 neuronal cluster labels, subsequently combined into an *M* x 20 matrix. Here, *M* denotes the trial count, and each row signifies the sequence of visited clusters during the respective trial.

### Trial similarity

To assess trial dynamics similarity, our focus was on highlighting the sequence of neuronal clusters. To achieve this, we designated the label −1 to each timepoint associated with the central point cloud. Meanwhile, other timepoints were allocated labels ranging from 1 to 7, corresponding to the respective neuronal cluster assignments. Subsequently, we created a trial structure matrix as previously outlined and computed pairwise trial similarity using cosine distance.

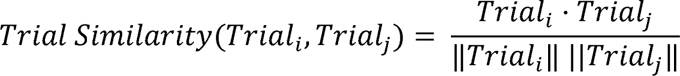

Here, *Trail_i_* represents the timepoint sequence of labels in the *ith* trial. This procedure resulted in a *M* x *M* for each dataset. Following that, we concentrated on the lower triangle of the matrix and pooled all values across all datasets.

### Decoding trial outcome across datasets

To enable across-datasets decoding, we generated a trial dynamics matrix using the same labeling approach employed for assessing trial similarity. Our goal was to enable decoding between datasets, which necessitated an alignment process. For each dataset, we initiated alignment by constructing a transition probability matrix between all neuronal clusters and the central point cloud. This process yielded a 8x 8 probability matrix for each dataset, except for one dataset with 7 clusters that was excluded from this analysis. Following this, the probabilities underwent clustering through the ward agglomerative complete linkage hierarchical clustering method^89^. Using the hierarchical clustering structure, we derived a cluster order based on the similarity of their probabilities. This order served as a mask across datasets to allow translation of clustering labels among dataset pairs. Essentially, this step aimed to address the question: Given transition probabilities, which cluster in dataset *A* most closely aligns with a cluster in dataset *B*. With the labels clusters of dataset *B* aligned to dataset *A*, we proceeded to compute the pairwise trial similarity between each trial pair in both datasets. For each trial *a* in dataset *A*, the decoded class of trial outcome was determined by maximizing the sum of Trial Similarity values over trials *b* in dataset B:

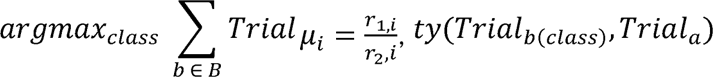

This process is analogous to identifying, on average after translation, which class of trials in dataset *B* most closely resembles an individual trial in dataset *A*. To further validate the decoding accuracy, we established a surrogate shuffle distribution by randomly shuffling trial classes in both datasets 200 times. We then calculated the decoding accuracy for each shuffle and computed the P-value by determining the percentile of shuffles that exhibited a higher decoding accuracy compared to the measured decoding accuracy. This entire process was repeated for every pairwise combination of datasets.

### Decoding trial outcome across datasets based on overall activity levels and thirsty-quenched linear axis

To decode trial outcomes across dataset based on overall activity levels, we calculated the overall activity levels within each trial by averaging the collective population activity across neurons. To decode trial outcomes across dataset based on the thirsty-quenched linear axis, we defined this axis for each mouse by 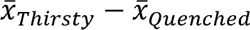, where 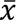 is the population vector of mean ongoing activity in each state. We projected peri-cue activity onto this axis by calculating the dot product of this vector with the time-varying pattern of InsCtx population activity, x(t). We then scaled values along this axis per mouse, ascribing a value of 1 when 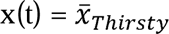, a value of 0 when 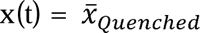 as 0, and intermediate values for patterns that fall between 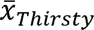 and 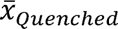 projected this data onto a thirsty-quenched linear axis, as outlined in reference^55^. Second, we utilized a similar decoding as explained above. Instead of training it with aligned trial labels for pairs of trials, we evaluated used the overall activity levels within each trial or the projection of the trial onto the thirsty-quenched axis as a training dataset.

### Quantification of anticipatory licking onset

The onset of anticipatory licking during a trial was defined as the time point when the first licking bout took place within the response window after cue presentation. Licking bouts were considered as instances where mice notably increased their licking rate. To detect these bouts, for each trial, we calculated the time intervals between consecutive licks occurring after cue presentation. We then identified the pair of licks with intervals shorter than a specific threshold:

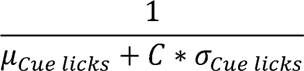

Where *μ_Cue licks_*, *σ_Cue licks_* were determined by calculating the average and standard deviation of licks based on the first 15 frames (0.5 sec) of cue presentation across all trials. The constant varied; it was set as 0.1 for datasets involving water rewards and 0.35 for datasets involving food rewards. The time point at which the first pair of licks exceeded this threshold was considered the onset of anticipatory licking.

### Quantification of cluster onset

To identify the onset of the cluster sequence, we first determined the two neuronal clusters with the highest occupancy during the response window for each dataset. Following this, we identified the initial time point within each trial where a transition occurred from the central point cloud to each of the identified clusters. The onset was defined as the first time point in each trial where the transition occurred with the cluster that exhibited the highest onset correlations with the lick onsets across all trials.

### Lick similarity between pairs of trials

To define the lick similarity between pairs of trials, for each trial, we smoothed the binary licks vector *Licks_i_*. To do so we applied a sliding window average of 100 frames (∼3.3 sec). The smoothed lick pattern for the *i*th trial was calculated as follows:

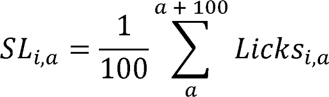

Where *a* is the *a*th licking frame in the original binary vector and *SL_i_* is the overall smoothed licking pattern. Lick similarity between the *i*th and *j*th trials was determined using the formula:

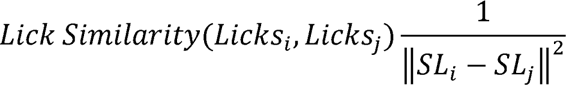

### Construction pseudo-trials in freely consuming mice

To construct pseudo-trials of consumption in freely consuming mice, we identified time points in which mice initiated a licking bout. To do so we identified all the time points within the free consumption epoch of the experiment in which the interval between two consecutive licks was above 1 second, unless this did not yield enough pseudo-trials, in which case we used 0.5 sec as the threshold (2/5 datasets). Every pair of licks that passed these criteria could be used to define the onset of a pseudo-trial. We then took all the time points that occurred two seconds before and four seconds after the second lick in that interval. We used these time points to identify the neuronal cluster dynamics and licking pattern that took place in the experiment and constructed a pseudo-trial matrix. We then removed pseudo-trials that overlapped with each other to avoid inflation of the matrix. To control for behavioral differences between operant task licking and free consumption licking, we repeated the analyses in (Fig. 4j-i) while selecting only pseudo-trials of free consumption in which the average licking rate in the first two seconds of the lick bout were within the mean ± standard deviation of the average licking rate in the first two seconds of operant task licking (Supp. Fig. 8c-e).

### Quantification and statistical analysis

All statistical details, including the specific statistical tests, are outlined in the corresponding figure legends. For unrelated samples from two different groups, we conducted a Wilcoxon rank sum test. In the case of matched-pairs related samples, we performed a Wilcoxon signed rank test if the number of samples exceeded five; otherwise, we used a matched pairs t-test. Dunn test was employed for multiple samples, encompassing all possible combinations of comparisons. When comparing distributions, such as the distribution of pairwise trial similarity, we employed a two-sample Kolmogorov-Smirnov test. A one-sided Pearson correlation moment test was carried out to determine the significance levels of sample correlations. To assess the significance levels of pairwise decoding, p-values were computed by extracting the percentile of the original decoding accuracy within the distribution of accuracies obtained by shuffling labels. For evaluating significance levels between pairs of manifolds, permutation tests were performed. The significance threshold was set at alpha = 0.05, and we corrected for multiple comparisons using Bonferroni-Holm correction.

### Declaration of generative AI and AI-assisted technologies in the writing process

During the preparation of this work the authors used Grammarly and ChatGPT in order to detect grammatical errors. After using these tools, the authors reviewed and edited the content as needed and take full responsibility for the content of the publication.

---

**<u>Video S1 – Stereotyped activity dynamics during rewarded trials on a single plane (mouse 1)</u>**

*Right*: Single plane of an example clustered structure. *Left*: Cluster dynamics heatmap in rewarded trials. Highlighted row on the cluster dynamics heatmap indicates the current rewarded trial, red dot on the clustered structure indicates current timepoint within the highlighted trial. Each timepoint is a 0.5 sec time-bin.

**<u>Video S2 – Stereotyped activity dynamics during rewarded trials on all planes (mouse 2)</u>**

*Right*: All planes of an example structure. *Middle*: All planes of the same clustered structure. *Left*: Cluster dynamics heatmap in rewarded trials. Highlighted row on the cluster dynamics heatmap indicates the current rewarded trial, red dots on the structure planes and clustered structure planes indicate current timepoint within the highlighted trial. Each timepoint is a 0.5 sec time-bin.

