## Supplemental figure and text for "Stereotyped goal-directed manifold dynamics in the insular cortex"

### **Supplemental Table 1**

| Figures | Experimental condition | Brain region | Data source | Number of mice | Number of datasets | Neurons per mouse (range) |
|---|---|---|---|---|---|---|
| **1 - 4, S1 - S7** | Operant task for water, Thirsty-Quenched | InsCtx | Livneh et al., 2020; unpublished data | 5 | 14 | 170-405 |
| **1, 3, 4, S1, S2, S3, S7, S8** | Operant task for Ensure, Hungry-Sated | InsCtx | Livneh et al., 2017; unpublished data | 7 | 10 | 59-247 |
| **1, 3, S2, S3, S7** | Operant task for Ensure, Hungry-Sated | Visual areas (V1, postrhinal) | Burgess et al., 2016 | 6 | 7 | 22-95 |
| **3, S7** | Operant task for Ensure, Hungry-Sated -AgRP activation | InsCtx | Livneh et al., 2017 | 4 | 4 | 70-82 |
| **3, S7** | Operant task for water, Thirsty-Quenched - SFO activation | InsCtx | Livneh et al., 2020 | 4 | 4 | 56-184 |
| **3, S7** | Operant task for water, Thirsty-Quenched- Hypertonic saline injection | InsCtx | Unpublished data | 3 | 3 | 89-160 |
| **4, S8** | Free Ensure consumption | InsCtx | Unpublished data | 5 | 5 | 111-224 |

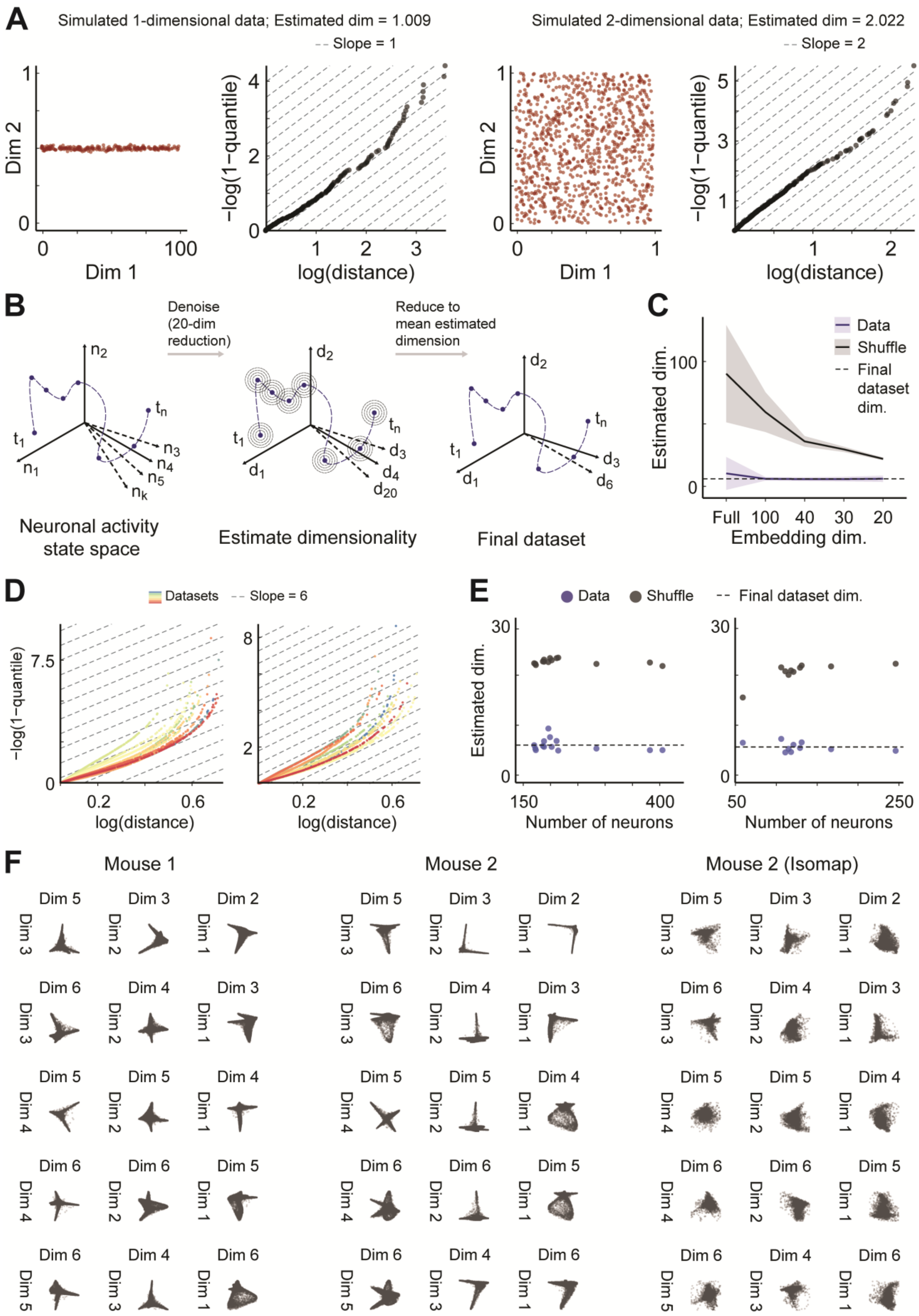
A
Simulated 1-dimensional data; Estimated dim = 1.009
Slope = 1
Simulated 2-dimensional data; Estimated dim = 2.022
Slope = 2
Dim 2
Dim 1
−log(1−quantile)
log(distance)
B
Denoise (20-dim reduction)
Reduce to mean estimated dimension
Neuronal activity state space
Estimate dimensionality
Final dataset
C
Data
Shuffle
Final dataset dim.
Estimated dim.
Embedding dim.
D
Datasets
Slope = 6
−log(1−quantile)
log(distance)
E
Data
Shuffle
Final dataset dim.
Estimated dim.
Number of neurons
F
Mouse 1
Mouse 2
Mouse 2 (Isomap)

**Figure S1 – Dimensionality estimation and reduction**

**A**. Dimensionality assessment using a two-NN Estimator on two simulated example datasets. Left: Estimation of a simulated one-dimensional dataset. Right: Estimation of a simulated two-dimensional dataset. In both datasets dimensionality is accurately approximated, with data points aligning parallel to the simulated dimension (i.e., slope).

**B**. Datasets initially undergo an intermediate dimensionality reduction, followed by an estimation of dimensionality for the intermediate dataset. The final dataset is achieved after an additional iteration of dimensionality reduction, using the mean estimated dimension in the intermediate iteration across all datasets.

**C**. Estimation of intrinsic dimensionality in shuffled and original data across different intermediate dimensionality reductions, and the full embedded data with no intermediate dimensionality reduction. Estimation of the intrinsic dimensionality in the original data converges to ~6 (dashed line, chosen for subsequent dimensionality reduction, see Figure S2). In contrast, estimation on shuffled data yields an intrinsic dimension close to the intermediate embedding dimension.

**D**. Left: Estimated dimensionality for 14 different datasets of thirsty mice. The near-constant slope observed across all datasets suggests a consistent intrinsic dimensionality of 6, which suggests the presence of approximately six latent variables within the activity manifold. Right: Same as in 'left' for 10 different datasets of hungry mice.

**E**. The estimated dimension was not correlated with the number of neurons sampled in each dataset for real data and shuffled data. Left and Right are the same as in '**D**' ($P > 0.2$, Pearson's product-moment correlation).

**F**. All dimension pairs in two different mice. Mouse 2 is presented with two different non-linear dimensionality reduction algorithms (right: Isomap; left: Laplacian eigenmaps, used throughout our analyses).

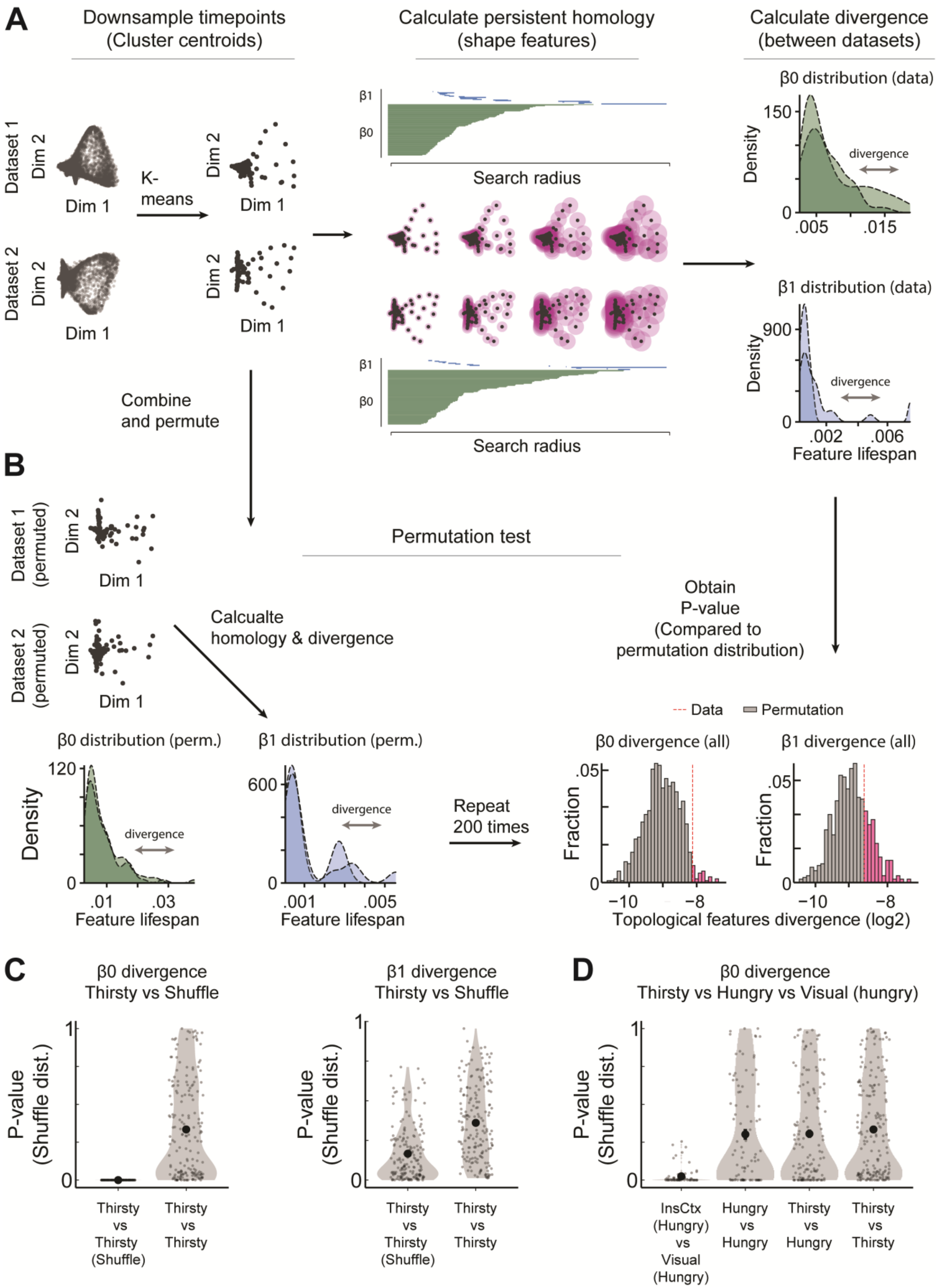
A
Downsample timepoints (Cluster centroids)
Calculate persistent homology (shape features)
Calculate divergence (between datasets)
Dataset 1
Dataset 2
Dim 1
Dim 2
K-means
β1
β0
Search radius
β0 distribution (data)
β1 distribution (data)
Density
divergence
Feature lifespan
Combine and permute
B
Dataset 1 (permuted)
Dataset 2 (permuted)
Permutation test
Calcualte homology & divergence
Obtain P-value (Compared to permutation distribution)
β0 distribution (perm.)
β1 distribution (perm.)
Repeat 200 times
Data
Permutation
β0 divergence (all)
β1 divergence (all)
Fraction
Topological features divergence (log2)
C
β0 divergence Thirsty vs Shuffle
β1 divergence Thirsty vs Shuffle
P-value (Shuffle dist.)
Thirsty vs Thirsty (Shuffle)
Thirsty vs Thirsty
D
β0 divergence Thirsty vs Hungry vs Visual (hungry)
InsCtx (Hungry) vs Visual (Hungry)
Hungry vs Hungry
Thirsty vs Hungry
Thirsty vs Thirsty

**Figure S2 - Assessing similarity between datasets using topological data analysis**

**A**. Illustration of the procedure used to assess the similarity between pairs of datasets. Initially, timepoints are down-sampled through K-means clustering. Subsequently, the distribution of topological features is extracted independently for each dataset. The magnitude of divergence in the feature distributions directly reflects the dissimilarity between the datasets, providing a quantitative measure of their dissimilarity.

**B**. Description the process for conducting permutation tests to assess the statistical significance of dissimilarity between datasets. Datasets are combined and permuted and redivided 200 times. During each permutation, the divergence of topological features between the randomly permuted and divided datasets is calculated. Subsequently, a p-value is derived by comparing the divergence observed between the original datasets to the distribution of divergences constructed from the permutations. This allows for the assessment of the statistical significance of dataset dissimilarity.

**C**. P-values calculated based on the shuffle distribution for all pairwise comparisons assessing the topological divergence of $\beta 0$ features.

**D**. Same as in '**C**' but for $\beta 1$ features. Shuffle distributions compared to neuronal data had non-significantly different results. This observation may be attributed to the sparse nature of the $\beta 1$ features extracted from each dataset.

**C**-**D**. N=100 pairwise comparisons for hungry vs hungry, N=182 pairwise comparisons for thirsty vs thirsty, N=140 pairwise comparisons for thirsty vs hungry mice, N=70 comparisons for Visual vs InsCtx mice, from N=14 datasets of thirsty mice, N=10 datasets of hungry mice, N=7 datasets of visual mice.

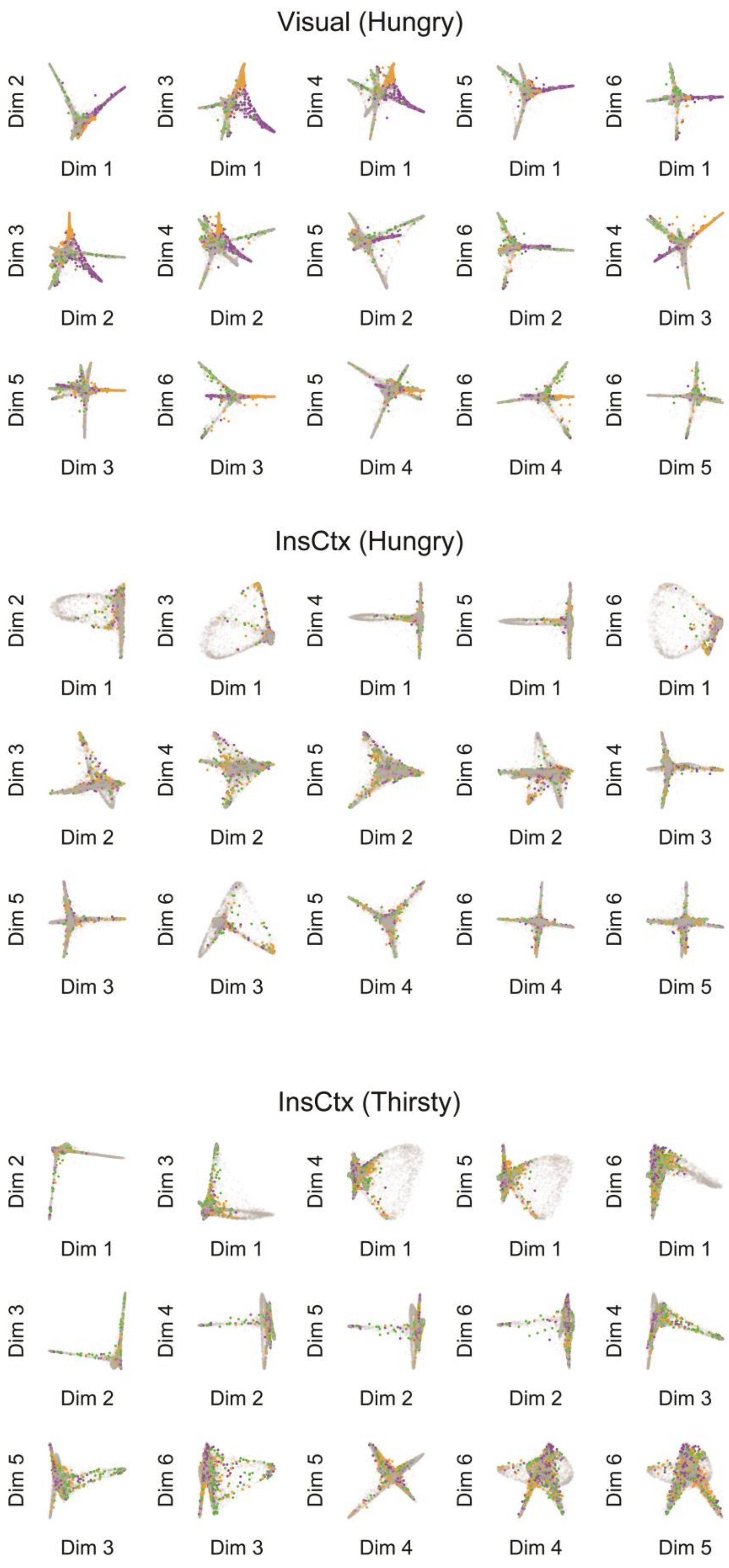

Visual (Hungry)
Dim 2
Dim 1
Dim 3
Dim 1
Dim 4
Dim 1
Dim 5
Dim 1
Dim 6
Dim 1
Dim 3
Dim 2
Dim 4
Dim 2
Dim 5
Dim 2
Dim 6
Dim 2
Dim 4
Dim 3
Dim 5
Dim 3
Dim 6
Dim 3
Dim 5
Dim 4
Dim 6
Dim 4
Dim 6
Dim 5
InsCtx (Hungry)
Dim 2
Dim 1
Dim 3
Dim 1
Dim 4
Dim 1
Dim 5
Dim 1
Dim 6
Dim 1
Dim 3
Dim 2
Dim 4
Dim 2
Dim 5
Dim 2
Dim 6
Dim 2
Dim 4
Dim 3
Dim 5
Dim 3
Dim 6
Dim 3
Dim 5
Dim 4
Dim 6
Dim 4
Dim 6
Dim 5
InsCtx (Thirsty)
Dim 2
Dim 1
Dim 3
Dim 1
Dim 4
Dim 1
Dim 5
Dim 1
Dim 6
Dim 1
Dim 3
Dim 2
Dim 4
Dim 2
Dim 5
Dim 2
Dim 6
Dim 2
Dim 4
Dim 3
Dim 5
Dim 3
Dim 6
Dim 3
Dim 5
Dim 4
Dim 6
Dim 4
Dim 6
Dim 5

**Figure S3 – Visualization of visual cortex and insular cortex neuronal manifolds with respect to visual cue presentation**

Visualization of the activity manifold in all dimension pairs of three example datasets. *Top*: visual cortex during hunger; *middle* and *bottom*: InsCtx during hunger and thirst, respectively. Data points are assigned three different colors (orange, green, purple) corresponding to timepoints during the three different visual cues (lasting two seconds). Note that timepoints associated with the three different visual cues appear segregated in the visual cortex manifold, but not in the InsCtx manifolds.

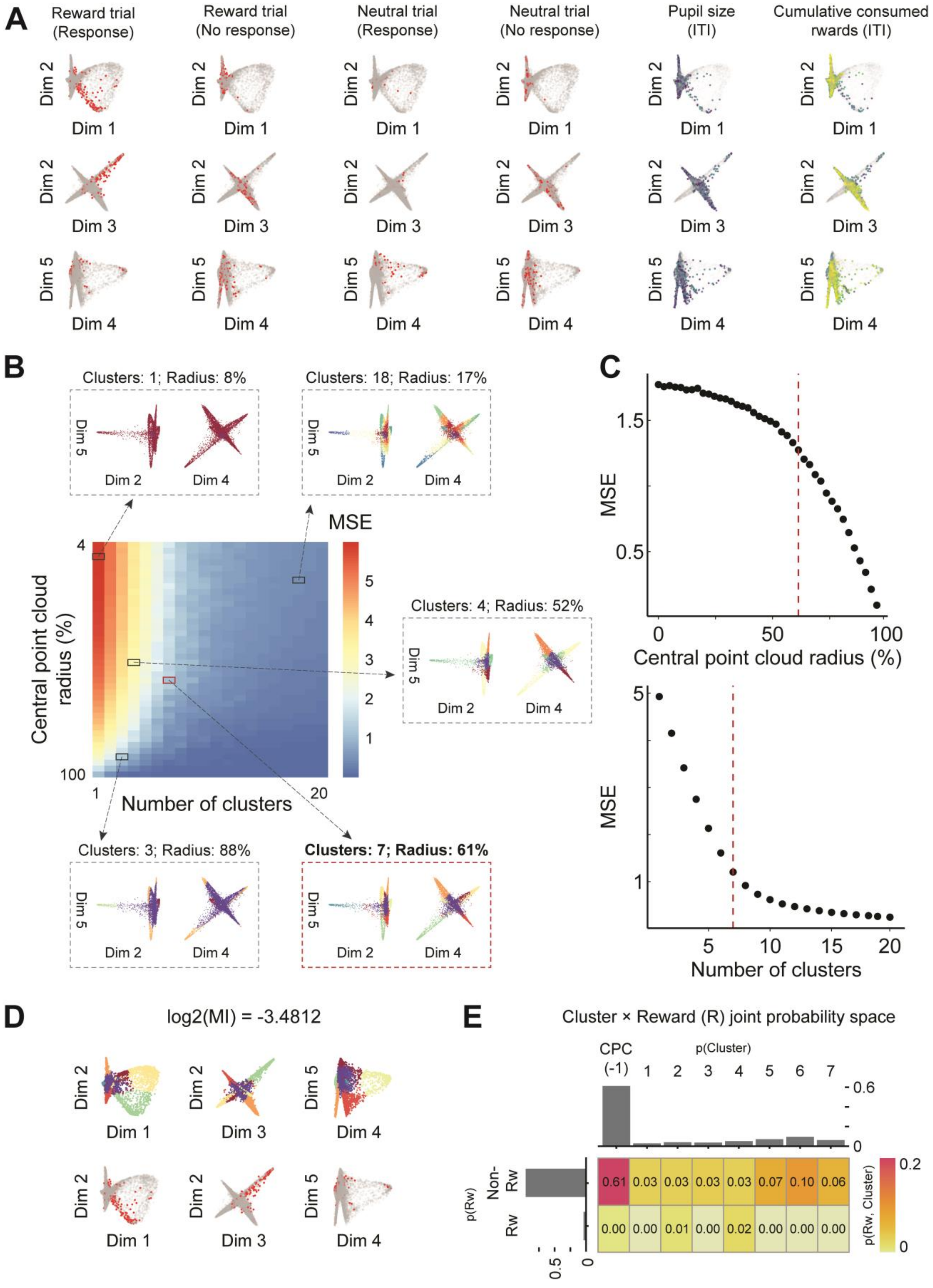
A
Reward trial (Response)
Reward trial (No response)
Neutral trial (Response)
Neutral trial (No response)
Pupil size (ITI)
Cumulative consumed rwards (ITI)
Dim 1
Dim 2
Dim 3
Dim 4
Dim 5
B
Clusters: 1; Radius: 8%
Clusters: 18; Radius: 17%
Clusters: 4; Radius: 52%
Clusters: 3; Radius: 88%
Clusters: 7; Radius: 61%
MSE
Central point cloud radius (%)
Number of clusters
C
MSE
Central point cloud radius (%)
MSE
Number of clusters
D
log2(MI) = -3.4812
E
Cluster × Reward (R) joint probability space
CPC (-1)
p(Cluster)
p(Rw)
Non-Rw
Rw
p(Rw, Cluster)
0.61
0.03
0.03
0.03
0.03
0.07
0.10
0.06
0.00
0.00
0.01
0.00
0.02
0.00
0.00
0.00

**Figure S4 – Automated clustering of the activity manifold; clustered structure relationship with experimental variables**

**A**. Example activity manifold annotated with external/internal variables. In each row, a distinct plane of the activity manifold is presented. To illustrate discrete variables (columns 1-4), we use red points to indicate the event occurrence. In the case of continuous variables (columns 5-6), points are color-coded, ranging from blue to yellow according to the variable's values.

**B**. Illustration of the automated clustering process. For every dataset, we construct a configuration matrix, as depicted. This matrix is generated by performing K-means clustering while iteratively adding points into the central point cloud (rows) and testing with varying cluster numbers (columns). The Mean Squared Error (MSE) is calculated for each K-means configuration during each repetition and stored in the corresponding matrix cell. Several examples of different configurations are shown in dashed gray boxes, whereas the final selected configuration is shown in a dashed red rectangle (see '**C**' for details).

**C**. Identification of the optimal configuration. The optimal number of points to include in the central point cloud and the optimal cluster count are automatically determined by identifying the deflection point, as detailed in the methods section.

**D**. Example planes of the clustered structure alongside an external variable of interest

**E**. Illustration of calculation of Mutual Information. The Mutual Information between the clustered structure and the external variable is obtained by calculating their joint probability space.

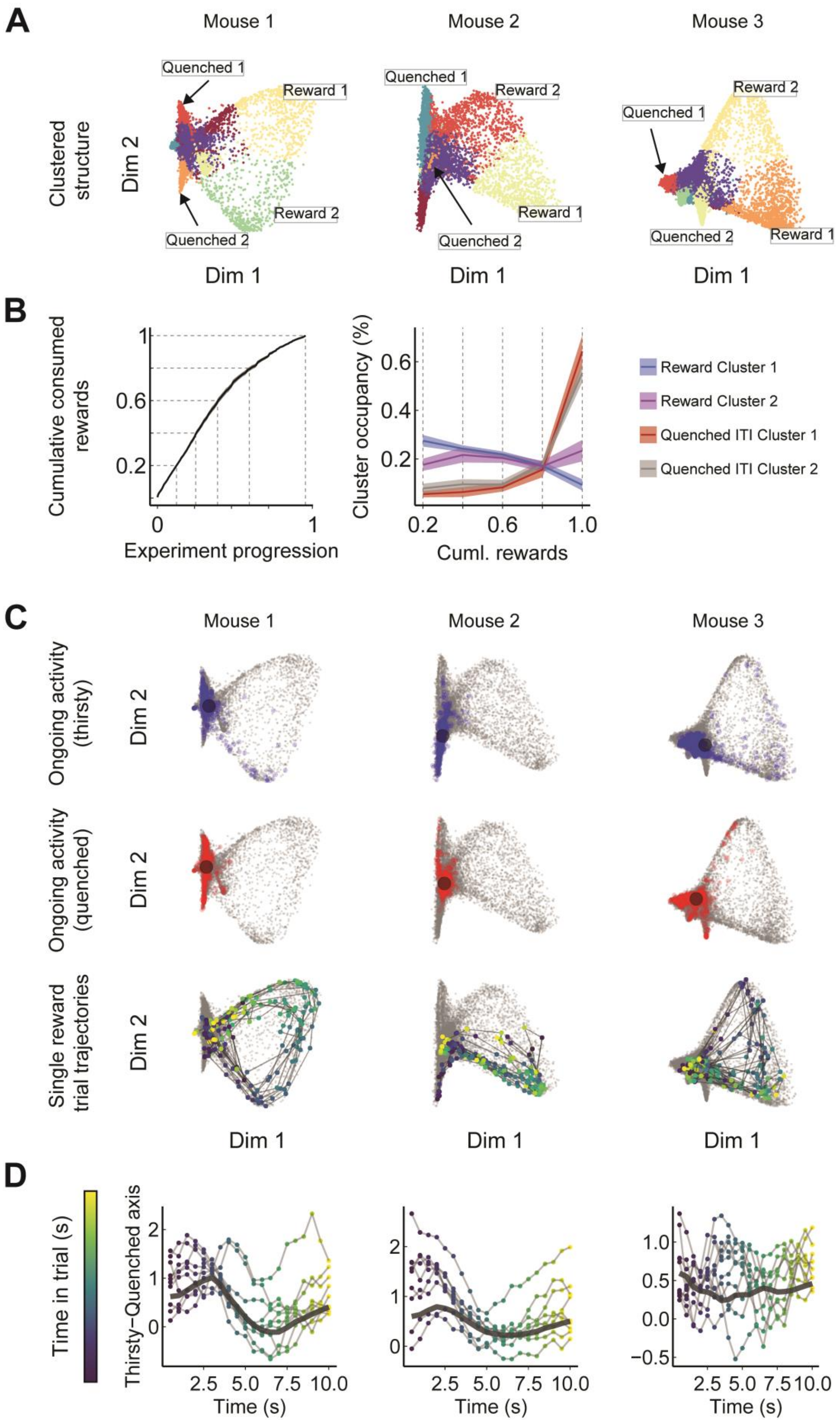
A
Mouse 1
Mouse 2
Mouse 3
Clustered structure
Dim 2
Quenched 1
Reward 1
Reward 2
Quenched 2
Dim 1
B
Cumulative consumed rewards
Experiment progression
Cluster occupancy (%)
Cuml. rewards
Reward Cluster 1
Reward Cluster 2
Quenched ITI Cluster 1
Quenched ITI Cluster 2
C
Ongoing activity (thirsty)
Ongoing activity (quenched)
Single reward trial trajectories
D
Time in trial (s)
Thirsty–Quenched axis
Time (s)

**Figure S5 – Comparison of linear projection dynamics with activity manifold dynamics**

**A**. Activity manifold from three example mice. Top: Neuronal clusters annotated using automated clustering, with labeled clusters representing ITI water satiety and reward clusters used in '**B**'.

**B**. Left: Physiological state approximated by cumulative consumed water rewards over time (mean across all datasets). Dashed lines indicate the experiment time at which mice reached a certain quenched level (e.g., by the middle of the experiment, mice consumed 70% of the rewards they would consume). Right: neuronal cluster occupancy (percent of time that activity occupied a certain cluster) during reward epochs and quenched phases in relation to gradual changes in physiological state (corresponding to dashed lines in the left cumulative rewards plot). Occupancy of reward clusters remained relatively consistent throughout gradually changing physiological states, and slightly decreased as the number of responded reward trials decreased with water satiation. In contrast, the two most frequently occupied neuronal clusters during the quenched phase were primarily visited when animals ceased water consumption and became quenched, but not before.

**C**. Top, middle: Timepoints of ongoing activity used to calculate average activity vectors for thirst (blue) and quenched (red) states, indicated by black centroids (following our previously published procedures[55]). Bottom: Trajectory of ten reward trials projected on the activity manifold.

**D**. The same ten reward trials as in '**C**', projected linearly using a neural mode obtained from subtracting the average ongoing activity in thirst and quenched states[55]. During rewarded trials, population activity briefly shifts from a thirst state towards a quenched state. The trajectory of the same trials can also be observed with respect to the activity manifold described in '**C**'.

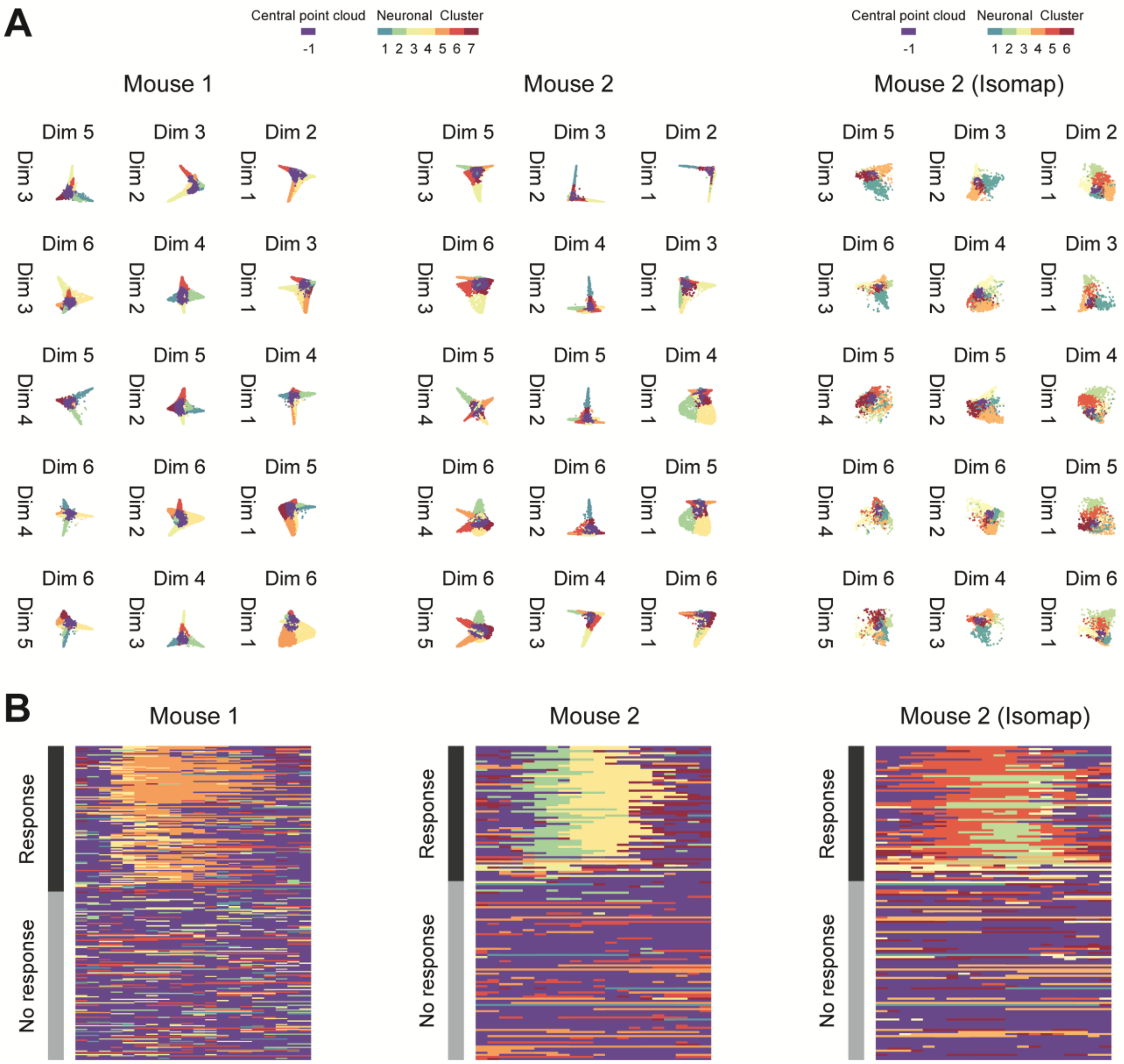
A
Central point cloud
Neuronal Cluster
-1
1 2 3 4 5 6 7
Central point cloud
Neuronal Cluster
-1
1 2 3 4 5 6
Mouse 1
Mouse 2
Mouse 2 (Isomap)
Dim 1
Dim 2
Dim 3
Dim 4
Dim 5
Dim 6
B
Mouse 1
Mouse 2
Mouse 2 (Isomap)
Response
No response

**Figure S6 – Activity dynamics of manifold cluster sequences with two different dimensionality reduction methods**

**A**. All dimension pairs in two different mice annotated by the automated clustering method. Mouse 2 is presented with two different non-linear dimensionality reduction algorithms (Right: Isomap; Left: Laplacian eigenmaps, used throughout our analyses).

**B.** Activity dynamics of the same examples as in '**A**' for reward cue trials. Notice that the stereotyped activity patterns are also observed when using a different non-linear dimensionality reduction algorithm (Isomap).

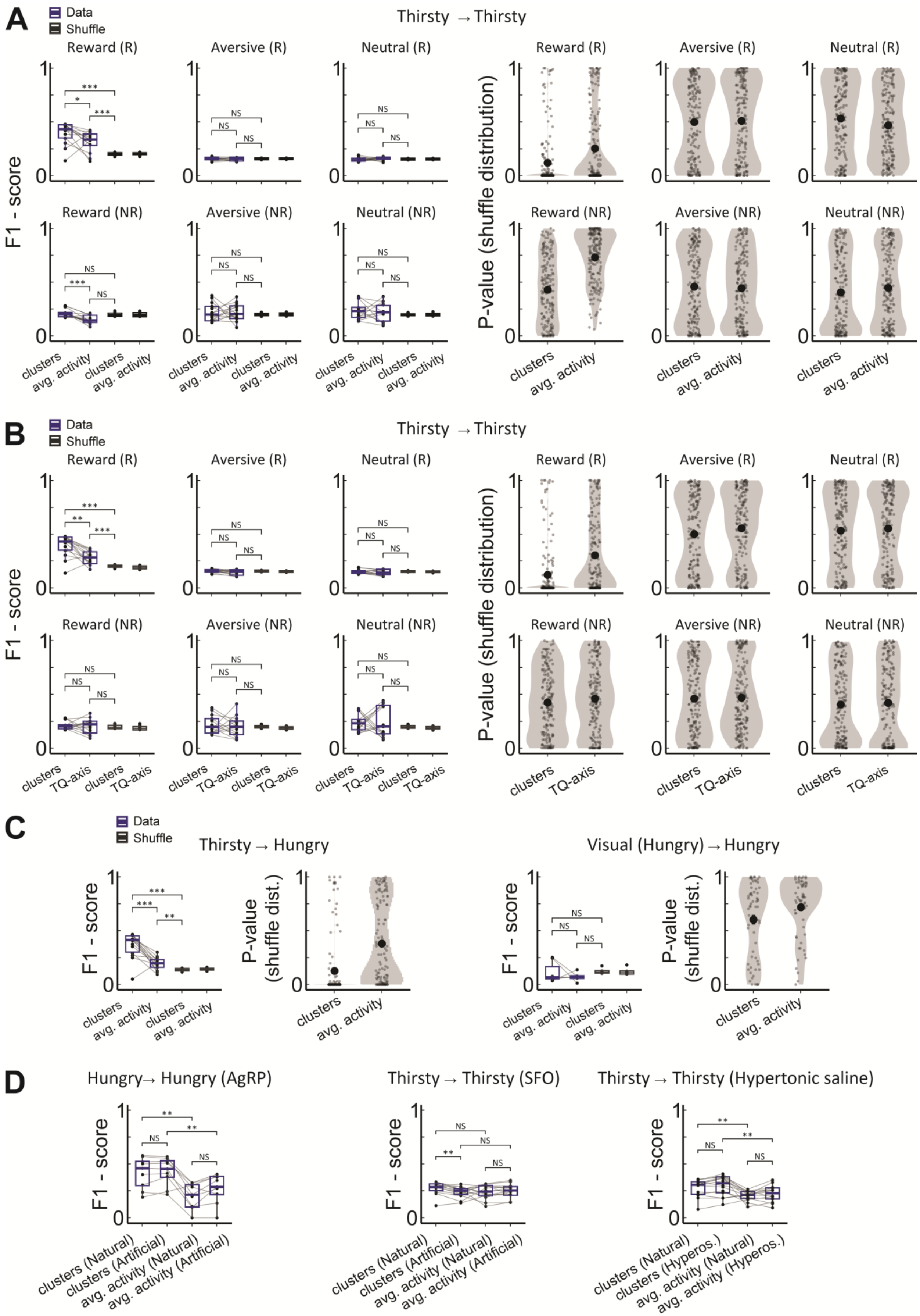

A
Data
Shuffle
Thirsty → Thirsty
Reward (R)
Aversive (R)
Neutral (R)
Reward (NR)
Aversive (NR)
Neutral (NR)
F1 - score
P-value (shuffle distribution)
clusters
avg. activity
NS
*
***
B
Data
Shuffle
Thirsty → Thirsty
Reward (R)
Aversive (R)
Neutral (R)
Reward (NR)
Aversive (NR)
Neutral (NR)
F1 - score
P-value (shuffle distribution)
clusters
TQ-axis
**
C
Data
Shuffle
Thirsty → Hungry
Visual (Hungry) → Hungry
F1 - score
P-value (shuffle dist.)
clusters
avg. activity
D
Hungry→ Hungry (AgRP)
Thirsty → Thirsty (SFO)
Thirsty → Thirsty (Hypertonic saline)
F1 - score
clusters (Natural)
clusters (Artificial)
avg. activity (Natural)
avg. activity (Artificial)
clusters (Hyperos.)
avg. activity (Hyperos.)

**Figure S7 – Analysis of across-dataset decoder performance**

**A.** Left: Assessing decoder F1-scores for trial types. A comparison of F1-scores when employing a decoder trained on manifold cluster sequences and another on average activity levels. Notably, only rewarded trials exhibited significantly higher decoding performance above chance. Additionally, decoding using manifold clusters outperformed decoding based on average activity levels. Right: The distribution of P-values for decoding derived from the surrogate shuffle distribution for each pairwise decoding is shown. All trial types exhibited uniform p-values in terms of F1-score significance levels, except for rewarded trials indicating significant decoding.

**B**. Same as in '**A**' but now comparing the performance of a decoder trained on manifold cluster sequences versus one trained on the projection of trials onto thirsty-quenched axes (T-Q axis).

**C**. Same as in '**A**', but now comparing between a decoder trained on InsCtx thirsty or visual cortex hungry datasets, and tested on InsCtx hungry datasets accordingly.

**D**. Comparison of the F1-score for rewarded trials across natural and artificial motivational states.

**A-D**. N=14 thirsty datasets, N=10 hungry datasets, N=7 visual datasets, N=4 AgRP datasets, N=4 SFO datasets, N=3 hyperosmotic datasets. $^*P < 0.05$, $^{**}P < 0.01$, $^{***}P < 0.001$, one-tailed Wilcoxon signed-rank test.

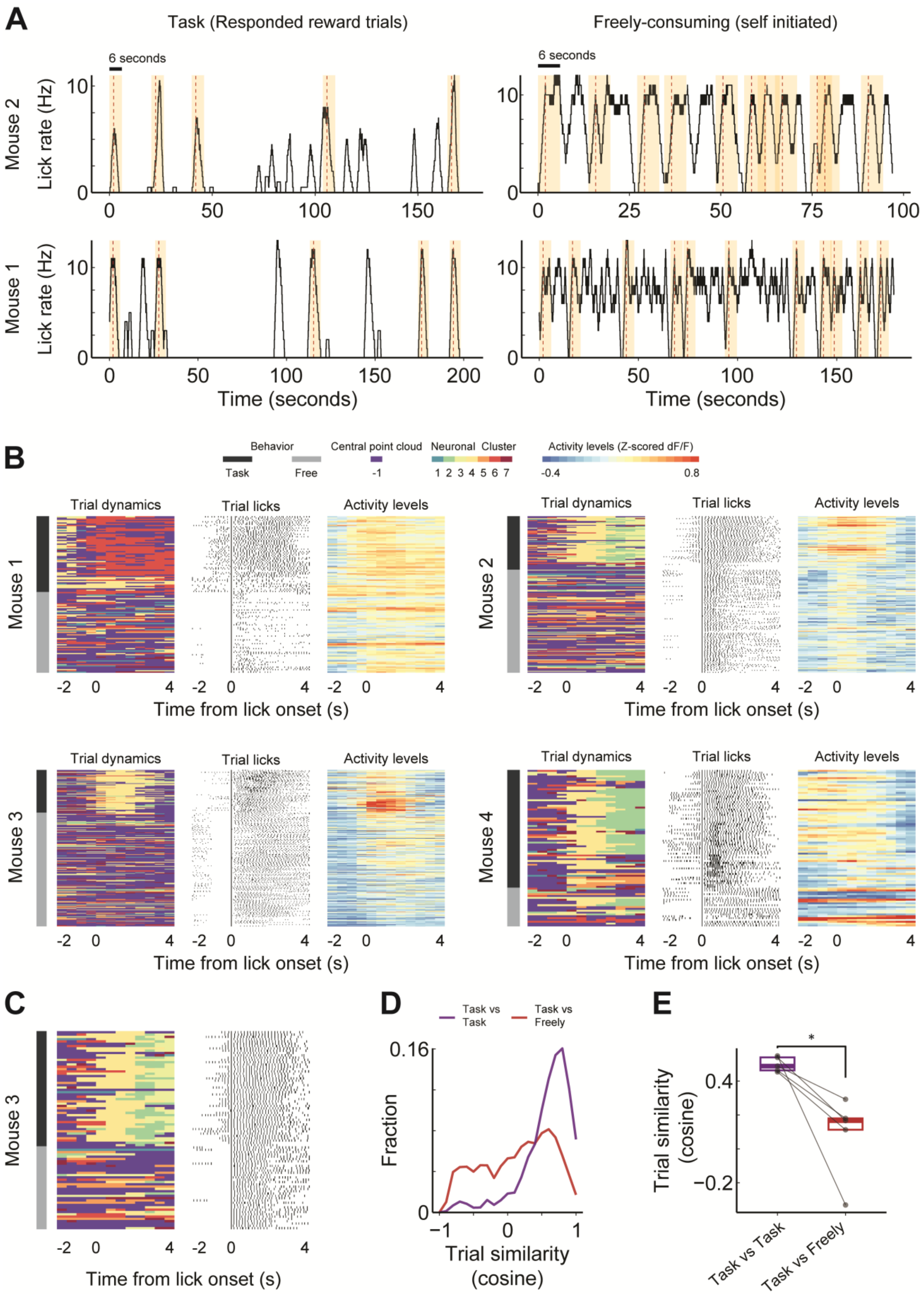
A
Task (Responded reward trials)
Freely-consuming (self initiated)
6 seconds
Mouse 2
Mouse 1
Lick rate (Hz)
Time (seconds)
B
Behavior
Task
Free
Central point cloud
-1
Neuronal Cluster
1 2 3 4 5 6 7
Activity levels (Z-scored dF/F)
-0.4
0.8
Trial dynamics
Trial licks
Activity levels
Mouse 3
Mouse 4
Time from lick onset (s)
C
D
Task vs Task
Task vs Freely
Fraction
Trial similarity (cosine)
E
*
Task vs Task
Task vs Freely

**Figure S8 - Comparison of activity dynamics in the goal-directed operant task vs. free consumption of rewards**

**A**. Examples of the pseudo-trial construction process to compare reward consumption behavior in two mice. Left: In the operant visual discrimination task, trials were generated by analyzing the 2 seconds before and the 4 seconds following lick onset in rewarded trials. Right: In free reward consumption, trials were generated by analyzing the 2 seconds before and 4 seconds after self-initiated lick bouts (timepoints in which licking rate sharply increased after a sufficient decrease). Only non-overlapping trials were chosen.

**B**. Example of activity dynamics during goal-directed behavior (operant visual discrimination task) and free reward consumption (self-initiated) in four different mice. Heatmaps are aligned with the constructed lick trials as described above. Left: Sequence of cluster dynamics observed in constructed lick trials. Middle: Licking in the constructed lick trials. Right: Overall mean activity levels. While goal-directed and self-initiated trials exhibit distinct dynamics, there is an increase in overall activity levels during both goal-directed and self-initiated consumption of rewards. Moreover, activity level changes on individual trials are not related to manifold dynamics.

**C**. Example cluster sequence dynamics and licking behavior, while controlling for variations in licking behavior between task-based consumption and freely initiated consumption. We addressed these behavioral distinctions by specifically choosing free-consumption trials in which the licking rate within the initial two seconds of the licking onset fell within the mean ± 1 standard deviation of the licking rate within the first two seconds of task-based lick bouts.

**D**. Distribution of all pairwise trial dynamics comparisons between reward consumption trials during task engagement and free consumption behavior, when controlling for behavioral differences in licking (pooled from N=5 datasets from mice).

**E**. Comparison of the average trial similarity for each dataset individually when comparing task dynamics to task dynamics, and task dynamics versus freely behaving consumption dynamics, while controlling for behavioral differences in consumption ($P < 0.05$, one-tailed paired t-test)
